# Dysregulation of the actin cytoskeleton and FMRP causes polar body protrusion defects in human fragile X premutation and aged oocytes

**DOI:** 10.1101/2025.02.23.639319

**Authors:** Kelly McCarter, Madhura Deshpande, Phillip Romanski, Christina Anna Stratopoulou, Nikica Zaninovic, Momina Tareen, Razan Al-Mousawi, Daylon James, Advaitha Madireddy, Zev Rosenwaks, Jeannine Gerhardt

## Abstract

Women carrying the fragile X premutation (55-200 CGG repeat expansion, PM) are at risk for developing fragile X-associated primary ovarian insufficiency (FXPOI), which is preceded by fragile X-associated diminished ovarian reserve (FXDOR). So far, the cause of FXDOR/FXPOI could not be comprehensively examined due to the scarcity of human ovarian tissue and oocytes. From studies in model systems, it was proposed that molecular abnormalities within the ovaries or a diminished primordial follicle pool cause FXDOR/FXPOI. To elucidate the defects instigating FXDOR/FXPOI, we examined human oocytes obtained from PM carriers undergoing *in vitro* fertilization (IVF). We found that the number of MII oocytes was reduced suggesting that the maturation of the oocytes is constrained in PM carriers. Furthermore, immature PM oocytes contained abnormal inclusions, irregular ubiquitin levels and DNA breaks. Despite these defects PM oocytes passed the DNA damage checkpoints. However, in anaphase I PM oocytes failed to initiate the protrusion of the first polar body. In addition, these oocytes amassed bundle actin structures, lacked an actin cap and had elevated profilin1 level. Profilin1 limits the formation of branched actin structures which are necessary for actin cap formation and membrane protrusions. Surprisingly, our results suggest that in PM oocytes an increase in FMRP elevates the profilin1 translation, which leads to the cytoskeleton defects and deficiencies in formation of the first polar body. We also analyzed the decline of MII oocytes in aging human ovaries. Similar, we found that the profilin1 expression and formation of the actin cytoskeleton were dysregulated due to appearance of cytoplasmatic FMRP foci in aged human oocytes. Thus, these results reveal that defects during anaphase I hinder the maturation of human oocytes resulting in FXDOR/FXPOI in PM carriers and a reduction in mature oocytes in women with advanced maternal age.

## Introduction

Diminished ovarian reserve and the decline of mature oocytes in patients with specific germline mutations and with advanced maternal age are not clearly understood. An expansion of a CGG repeat sequence within the 5’ untranslated region of the *FMR1* gene is associated with several disorders. While patients with more than 200 CGG repeats express the full mutation and the classic phenotype of Fragile X Syndrome (FXS), patients with 55-200 CGG repeats are premutation (PM) carriers and are at risk for fragile X-associated tremor/ataxia syndrome (FXTAS) and fragile X-associated primary ovarian insufficiency (FXPOI), which is preceded by fragile X-associated diminished ovarian reserve (FXDOR) (reviewed in (Hagerman and Hagerman, 2004; Man et al., 2017; Willemsen et al., 2011)). It is estimated that 1 in 150-300 women carry the PM (Hunter et al., 2014). Compared to the general population, PM carriers are approximately 16 to 20 times more likely to develop POI, which is characterized by menstrual irregularities and infertility (Allen et al., 2021). The increased risk of FXPOI and FXDOR is non-linear to the repeat number, with the mid-range repeat size (70-90 CGG repeats) having the greatest risk of FXDOR/FXPOI (Allen et al., 2007; Lekovich et al., 2018). However, the underlying defects and mechanisms causing of FXDOR/FXPOI in PM carriers remain unclear.

Mammalian oocytes are generated from female germ cells and arrested at prophase I, which reside in primordial follicles. Following puberty, the cyclic luteinizing hormone (LH) surge initiates resumption of meiosis I of the oocyte within the pre-ovulatory follicle, resulting in oocyte maturation. It was suggested that in PM carriers, the primordial follicle pool might be depleted, or the establishment of the pool is inhibited during prenatal development. Other considerations are that factors intrinsic to the ovary contribute to the development of FXDOR/FXPOI (Man et al., 2017). These mechanisms may have a detrimental effect on the ovary and therefore lead to increased rates of follicular atresia. In addition, from studies in model systems, different mechanisms were proposed as the cause of FXDOR/FXPOI. Using Drosophila as a model, it was found that FMRP is required for preservation of germline stem cells in the ovaries (Yang et al., 2007). In contrast, in a PM mouse model the primordial follicle pool was not affected, however, there was a reduction in the number of advanced subclasses of follicles (Hoffman et al., 2012). It was also reported that cells in model systems and in PM patients contain an increased amount of mutant FMR1 mRNA (which contain the repeat expansion) and a reduced number of FMRP protein (Kenneson et al., 2001; Tassone et al., 2000).

Additional commonly observed features in cells of PM patients are formation of FMR1 mRNA aggregates and ubiquitin-positive intranuclear inclusions (Galloway and Nelson, 2009). Furthermore, in brain tissues of PM patients with FXTAS, a neurological disorder affecting primarily males (Hagerman et al., 2001), abnormal FMR1 polyglycine (FMRpolyG) inclusions were found that are generated by RAN translation (Todd et al., 2013). FMRpolyG inclusions and FMR1 RNA aggregates were also detected in granulosa cells (Friedman-Gohas et al., 2020; Rosario et al., 2022). Granulosa cells are ovarian cells that surround the oocytes in the follicles and facilitate the oocyte maturation. Recently, expression of the FMRpolyG inclusions were detected in ovarian stromal cells, however, this one human ovary specimen that was studied did not contain any follicles or oocytes (Buijsen et al., 2016). Thus, it is still not clear if inclusions in stroma cells cause FXDOR/FXPOI or if human oocytes itself contain abnormal FMRpolyG inclusions and other defects that cause FXDOR/FXPOI.

To determine the mechanisms and defects leading to FXDOR/FXPOI, we analyzed human oocytes obtained from PM carriers going though IVF. We uncovered that PM patients, in particular those with mid-range repeat size, have a lower proportion of oocytes that are mature (MII oocytes) suggesting defects during the maturation of the PM oocytes. We also detected irregular ubiquitin staining and FMRpolyG inclusions. Despite these cellular abnormalities and high number of DNA breaks, PM oocytes pass the DNA damage checkpoints and progress into anaphase I. We found that spindle migration to the cortex and separation of the chromosomes was not disturbed. However, the formation of the actin cap and the extrusion of the polar body was defective PM oocytes. Furthermore, we detected an elevated level of profilin1, which is known to limit the branched actin structure formation needed for membrane protrusions. Profilin1 mRNA is a target of FMRP. In contrast to other PM cells, the translation of the mutant FMR1 mRNA containing the expanded CGG repeats was not disturbed in PM oocytes and surprisingly elevated. These results reveal that in PM oocytes an increase in FMRP expression activates disproportionate the profilin1 translation leading to dysregulation of the cytoskeleton and defects in protrusion of the first polar body. Thus, our data suggest that in PM oocytes dysregulation of FMRP protein cause defects that lead to the disease symptoms. Since the oocytes quantity is also reduced in women with advanced maternal age, we also study unaffected aged oocytes. We found similar to PM patients elevated profilin1 levels and defective actin cap formation in aged oocytes. However, FMRP was not elevated instead the protein was dislocated into the ooplasm. These results reveal that dysregulation of cytoskeleton structures and FMRP are accountable for the reduction of mature oocytes and FXDOR/FXPOI in PM carriers as well as contribute to the decline in oocytes quantity with advance maternal age.

## Results

### PM patients have less mature MII oocytes but similar number of GV and MI oocytes

It is not clear whether FXDOR/FXPOI is caused by a decrease in the number of primordial follicles, defects during oocyte maturation or oocytes degeneration by follicular atresia. To elucidate the mechanism and defects causing of FRDOR/FXPOI, we first examined which step during oocyte development and maturation was affected. Therefore, we performed a retrospective analysis of 34 PM carriers (55-200 repeats) who underwent 83 IVF cycles, 44 intermediate allele (IM, 45-54 repeats) carriers who underwent 96 IVF cycles and 207 control patients identified to have <45 CGG repeats in both FMR1 genes who underwent 249 cycles. **Table 1** and **Suppl. Table 1** display the demographic and baseline characteristics of the IVF patients. The mean age in the PM and control groups (< 45 repeats) was 33.8 years old (*p* = 0.9). The majority of the recipients had a normal BMI and had not previously given birth. The groups had similar gravidity, parity, and BMI (**Suppl. Table 1**). As previously reported (Lekovich et al., 2018) despite a similar age and BMI, PM group had a lower Anti-Mullerian Hormone level (AMH) compared to the control group (2.3 ng/ml vs 3.2 ng/ml; *p* = 0.02). The PM group also had previously undergone more IVF cycles than the control group (2.3 cycles vs 1.3 cycle; *p*=0.002). The mean age in the IM allele group was 37 years old, which was somewhat older than the PM and control groups. However, despite an older age, the AMH level was higher in the IM allele group compared to the PM group and were similar to the control group. **Table 1** and **Suppl. Table 2** display the IVF cycle outcomes. Peak estradiol level, total gonadotropin dosage and days of stimulation were similar between the groups. There were no significant differences between the percent of immature and mature oocytes (**Suppl. Table 2**). However, the PM group had a lower number of total oocytes retrieved (13.1 vs 15.9; *p*=0.02) compared to the control group as seen previously (Lekovich et al., 2018) and specifically a lower number of mature oocytes (10.4 vs. 12.3; *p*=0.05) compared to the control group.

**Table 1.** Demographic and baseline characteristics of IVF patients. The mean age, BMI, AMH and number of previous IVF cycle are displayed for control and PM carriers. The groups are similar for age and BMI. The number of retrieved oocytes per cycle, including mature MII oocytes, are also presented for control and PM carriers. The standard deviation and *p*-value are indicated.

|  | Control Group<br>(n=207)<sup>†</sup> | PM Group (n=34)<sup>†</sup><br>*p*-value* | IM Group (n=44)<br>*p*-value** |
| --- | --- | --- | --- |
| CGG repeat length | <45 CGG repeats | 55-85 CGG repeats | 45-54 CGG repeats |
| Patient age (years) ± SD | 33.8 ± 4.0 | 33.8 ± 3.9<br>0.9 | 36.8 ± 3.7<br><0.001 |
| Body Mass Index (kg/m<sup>2</sup>) ± SD | 22.1 ± 8.0 | 20.8 ± 7.7<br>0.2 | 23.7 ± 4.0<br>0.01 |
| Mean AMH (ng/ml) ± SD | 3.2 ± 3.2 | 2.3 ± 2.3<br>0.02 | 3.4 ± 2.9<br>0.4 |
| Mean Previous IVF cycle ± SD | 1.3 ± 1.6 | 2.3 ± 2.0<br>0,002 | 1.8 ± 2.8<br>0.1 |
| Mean Number of Oocytes Retrieved ± SD | 15.9 ± 9.5 | 13.1 ± 9.4<br>0.02 | 14.0 ± 9.6<br>0.1 |
| Mean Number of MII Oocytes ± SD | 12.3 ± 7.8 | 10.4 ± 7.8<br>0.05 | 10.5 ± 7.0<br>0.04 |
| <sup>†</sup> n refers to number of individual patients<br>* *p*-value compares PM group to control group<br>** *p*-value compares IM group to control group |  |  |  |

Next, to analyze which specific steps in oocyte development is affected in PM carriers, the number and percentages of immature and mature oocytes were compared to the number of CGG repeats. After egg retrieval the embryologists graded the oocytes by presence or absence of a polar body. Mature human oocytes have completed meiosis I and are arrested in metaphase II (MII) thus have a polar body present (**Fig. 1a**). Immature oocytes do not have a polar body and were further classified upon present or absence of the germinal vesicle (GV). Oocytes with GVs are classified as GV oocytes and if absent as MI oocytes. GV oocytes are arrested in prophase I. Next, we evaluated the association between repeat number and oocyte maturity. We found that the percent of MII oocytes decreased significantly with increasing repeat number (**Fig. 1b-d**). However, no major differences were found between percent of GV and MI oocytes in PM carriers.

**Figure 1.**
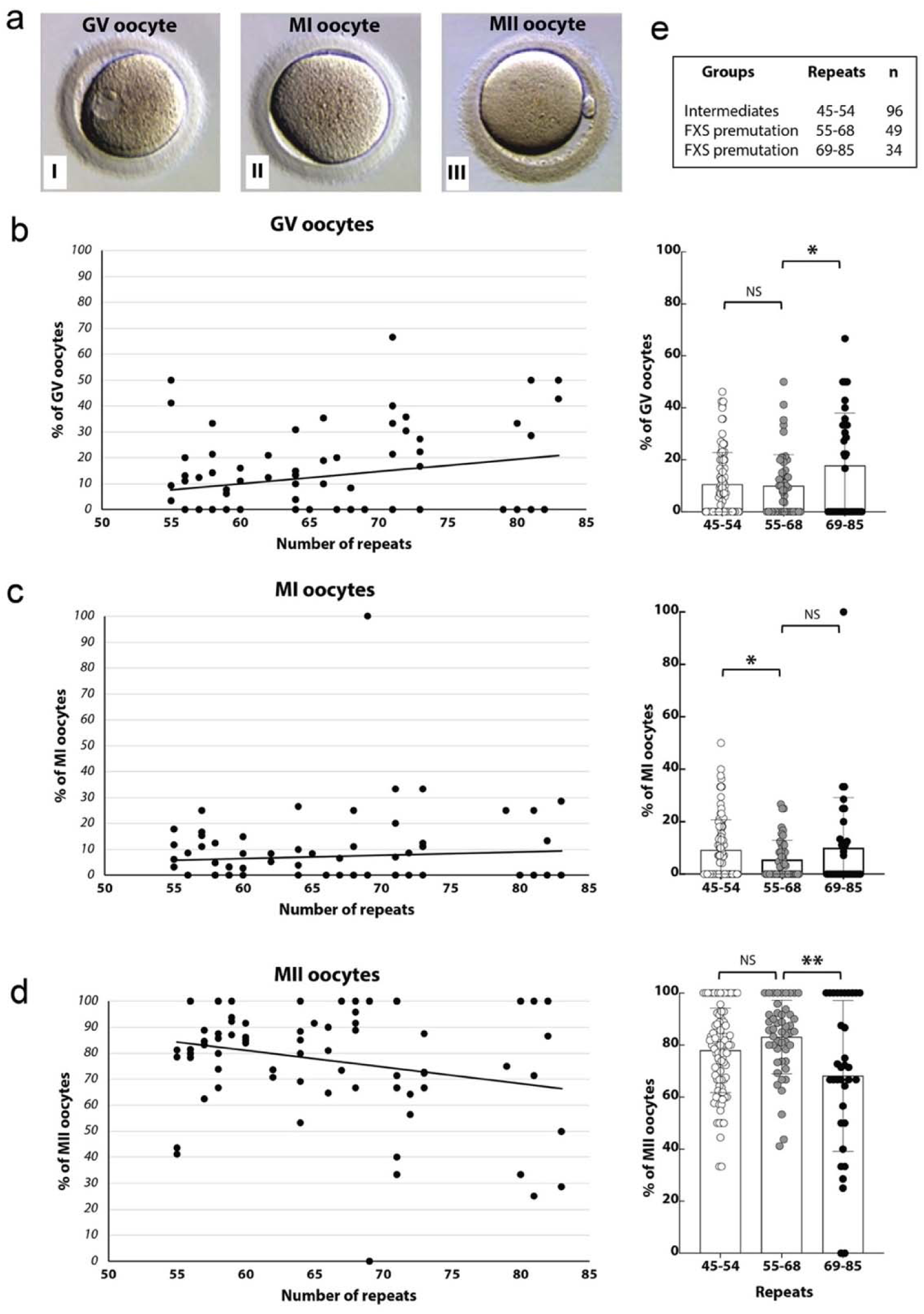
A reduced percentage of mature MII oocytes were detected in PM carriers. **a)** Grading of oocytes in immature GV (i) and MI (ii) oocytes without a polar body and mature MII (iii) oocytes that contained a polar body. **b)** Diagrams of the percent of GV (**b**), MI (**c**) and MII (**d**) oocytes are plotted over the CGG repeat length (55 to 85 CGG repeats). **e)** PM patients were divided into three groups based on repeat number (intermediate (45-54), PM from 55-68 and from 69-85 repeats). We divided the PM carriers in two groups, low repeat number PM (55-68 CGG) and mid-range repeat number PM (69-85 CGG) group. The number of cycles (n) is indicated in the table. Below the average of the percent of GV, MI and MII oocytes are shown. Standard deviation and *p*-values (NS, not significant, * < 0.05, ** <0.005) are indicated.

Since it was reported that PM patients with mid-range CGG repeat size have the highest risk for FXDOR/FXPOI, we compared the percentage of GV, MI and MII oocytes in PM carriers with low and mid-range CGG repeat lengths. The patients were divided into three groups based on repeat number (45-54, 55-68 and 69-85 repeats) (**Fig. 1e**). We included in our analysis intermediate carriers (IM) with 45-54 CGG repeats and divided the PM carriers into low repeat number (55-68 CGG) and mid-range repeat number (69-85 CGG) PM groups. We found that patients with 69-85 repeats had a significantly lower percentage of MII oocytes compared to the other two groups. Interestingly, the percent of GV oocytes increased slightly in patients with 69-85 repeats. There was no significant association between percent of MI oocytes and repeat number. Next, we analyzed the total number of GV, MI and MII oocytes (**Suppl. Fig. 1a-d**). We found that PM carriers with 69-85 repeats had a significantly lower number of mature MII oocytes than the other two groups. In summary, these results reveal that in PM carriers oocyte maturation is not perturbed in GV and MI phase, however there is a drop in the number of mature MII oocytes.

### PM oocytes display cellular abnormalities, including FMRpolyG inclusions and irregular ubiquitin pattern

To elucidate the cellular defects that lead to the lower proportion of MII oocytes in PM carriers, we analyzed the discarded immature PM oocytes, which we obtained from patients going through IVF. All control and PM oocytes examined in this study are immature, marked by the absence of the polar body. One of the most commonly observed features in brain tissue of PM patients with FXTAS are ubiquitin-positive intranuclear inclusions (Galloway and Nelson, 2009), which contain aberrant FMRpolyG peptides (Todd et al., 2013). It was suggested that these inclusions instigate cell dysfunction and the disease symptoms by sequestering proteins and causing the loss of their cellular functions. First, we analyzed the PM oocytes for ubiquitin inclusions and the presence of FMRpolyG aggregates by immunostaining (**Fig. 2a, Suppl. 2a**). In immature control and IM oocytes (**Suppl. Fig. 2a-b**) we detected no inclusions and no FMRpolyG foci. However, FMR1polyG foci were detected in PM oocytes (**Fig. 2a, Suppl. Fig. 2a**). In addition, we detected ubiquitin staining in all oocytes but no large inclusions and the FMRpolyG foci did not overlap with the ubiquitin staining. These results differ from the ubiquitin positive inclusions, which overlap with the abnormal FMRpolyG peptide in FXTAS brain tissue and other cell types of PM patients.

**Figure 2.**
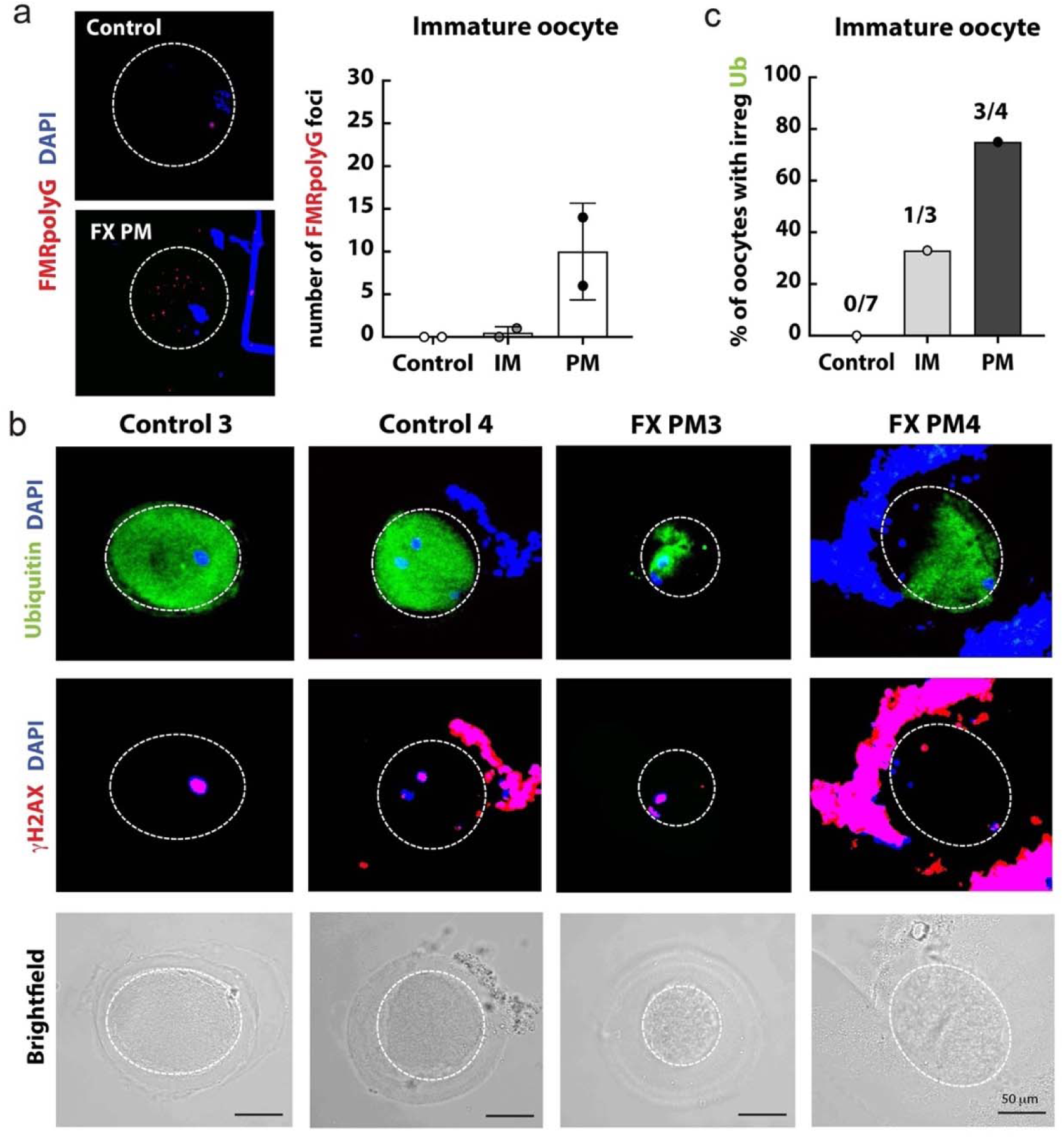
PM oocytes contain FMR1polyG aggregates, irregular ubiquitin staining and DNA breaks. **a) Left;** Analysis of immature control and PM oocytes stained with FMRpolyG (red) and ubiquitin antibody (green). The DNA was visualized with DAPI. **Right;** The FMR1polyG foci were quantified in human oocytes; two control, two PM and two intermediate (IM) oocytes (**Suppl. Fig. 2a-b**). **b**) Control and PM oocytes were stained with ubiquitin antibody (green), and γH2AX antibody (red) to visualize the DNA breaks. The DNA was detected with DAPI (blue). Some oocytes were not so well denuded and still contained the surrounding granulosa cells. **c**) Analysis of the ubiquitin staining in oocytes (control, IM and PM from **b** and **Suppl. Fig. 2** and **3**). IM oocytes are pictured in **Suppl. Fig. 3b**.

PM oocytes also had aberrant ubiquitin staining that was irregular and reduced in contrast to control oocytes (**Fig. 2b**). Ubiquitin is a small polypeptide that is involved in the regulation of protein degradation, cell cycle checkpoint, transcription and DNA repair. Previously cells from ALS patients, also a disorder marked by abnormal inclusions, displayed reduce monomeric ubiquitin (Farrawell et al., 2020). Analysis of additional control and PM oocytes confirmed the irregular ubiquitin staining in PM oocytes (**Fig. 2c** and **Suppl. Fig. 2a and 3a**). One IM oocytes also showed an aberrant ubiquitin pattern (**Suppl. Fig. 3b**). In contrast control oocytes showed even ubiquitin staining throughout the oocyte, including the immature oocytes that were damaged or started to degenerate (**Suppl. Fig. 3a**). In parallel, we stained these oocytes for γH2AX, a marker of double-strand DNA breaks (DSB, **Fig. 2b** and **Suppl. Fig. 3a-b**). All oocytes contained high level of DSB breaks. The finding of DNA breaks in these oocytes was not surprising since all oocytes are discarded immature or damaged oocytes obtained from patients going through IVF. The surrounding granulosa cells, which were still attached to some PM oocytes, also showed a high level of DNA breaks (**Fig. 2b**). However, granulosa cells had little to no ubiquitin staining. In summary, these findings reveal that PM oocytes contain specific cellular abnormalities, such as irregular ubiquitin pattern and aberrant FMRpolyG inclusions.

**Figure 3.**
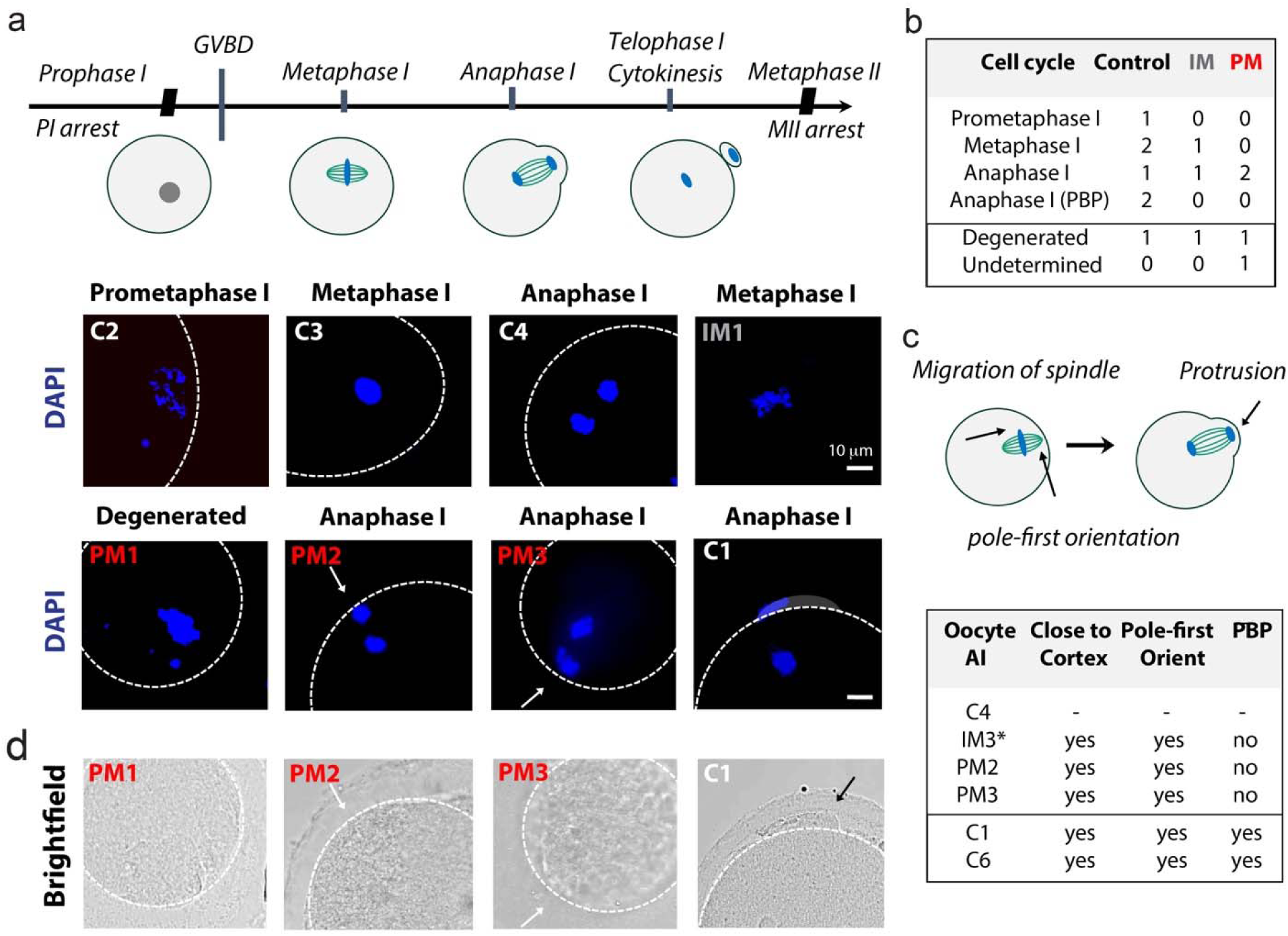
Defects during anaphase I inhibit the maturation of PM oocytes. **a) Top;** Schematic representation of the cell cycle phases of meiosis I. **Bottom**; DAPI staining of control, IM and PM oocytes. **b**) Table summarizes the number of oocytes in different the cell cycle phase and includes number of degenerated oocytes and oocytes where cell cycle phase could not be determined. **c**) **Top;** In metaphase I the spindle and DNA migrates to the oocyte cortex in pole-first direction. Then extrusion of the polar body is initiated concurrently with the separation of the homologous chromosome pairs. One set of the chromosomes are pulled into the protruding polar body (also see **a** and **d**, C1 and **Suppl. Fig. 4a**, C6). **Bottom**; Table is summarizing the characteristics of the anaphase I (AI) oocytes and if in these oocytes the spindle is accurately positioned at the cortex and protrusion of the polar body (PBP) is initiated. * In IM3 oocyte (**Suppl. Fig. 3b)** chromosome pairs just started to be separated, thus this oocyte might not initiate polar body (PB) protrusion yet. **d**) Brightfield pictures of three PM and one control oocytes, where protrusion of the polar body started (black arrow) or was absent (white arrow).

### PM immature oocytes pass the DNA damage checkpoint but do not complete meiosis I

Although some degree of DNA damage is tolerated and does not inhibit the maturation of oocytes, severe DNA damage triggers prophase I arrest at the G2/prophase DNA damage checkpoint, or subsequently oocytes arrest during metaphase I due to the activation of the Spindle Assembly Checkpoint (SAC). To determine if the immature PM oocytes arrest at these checkpoints, we examined the cell cycle phases and development stages of these oocytes in meiosis (**Fig. 3a**). DAPI staining of the genomic DNA of the control oocytes revealed that six of these oocytes were in different meiotic phase (prometaphase I, metaphase I, anaphase) and one oocyte was degenerated (**Fig. 3b** and **Suppl. 3a**). DAPI staining of the IM oocytes showed oocytes in metaphase I and anaphase I (**Fig. 3a** and **Suppl. 3b**). From the four PM oocytes, one oocyte was degenerated, and two PM oocytes were in anaphase I of meiosis I (**Fig. 3a** and **b**). From one PM oocyte we could not determine the cell cycle stage due to the surrounding and overlapping granulosa cells (**Fig. 2b**, right side). However, DAPI staining of the other PM oocytes showed that PM oocytes were able to mature to anaphase I. These results reveal that despite cellular abnormalities and high amount of DNA breaks, PM oocytes are able to pass the DNA damage checkpoints and enter anaphase I.

At the onset of meiosis I chromosomes condense, and microtubules reorganize around them into a bipolar spindle (Schuh and Ellenberg, 2007). In metaphase I, the spindle with the chromosomal DNA then migrates towards the oocyte periphery in a pole-first orientation so that a perpendicular orientation of the spindle is achieved at the cortex without rotation (Verlhac et al., 2000) (**Fig. 3c**, top). Migration occurs along the spindle axis and is directed towards the closest border of the oocyte. Once the spindle is translocated to the oocyte cortex, the homologous chromosome pairs are separated, travel poleward and one set of chromosome is pulled into the polar body in anaphase I (Fabritius et al., 2011). Protrusion of the first polar body is initiated at the same time as the separation of the chromosome pairs in anaphase I (Greaney et al., 2018; Wei et al., 2018). The polar body formation is then completed during telophase I. Analyzing the orientation and location of the chromosomal DNA, we noticed that in PM oocytes the spindle and DNA is located correctly at the cortex in pole-first orientation and the chromosomes are already separating and traveling towards the poles (**Fig. 3a** and **c**, bottom). However, the onset of polar body protrusion was not observed in PM oocytes in anaphase I as reported previously (Greaney et al., 2018; Wei et al., 2018) and seen in control oocytes (**Fig. 3d** and **Suppl. Fig. 4a**). In summary, PM oocytes are able to pass the DNA damage and spindle positioning checkpoints (Metchat et al., 2015) as well as show no defects in chromosome separation, but these oocytes failed to initiate polar body extrusion to complete meiosis I.

### Actin cap formation needed for polar body protrusion is defective in PM immature oocytes

Since we observed that the onset of polar body protrusion was lacking in PM oocytes, we were wondering next if the actin cap is formed in these oocytes. Mammalian oocytes contain a cortical actin layer at the periphery. After the spindle reaches the cortex the cortical actin directly overlying the spindle apparatus thickens, forming the actin cap (Longo and Chen, 1985) (**Fig. 4a**). The actin cap is essential for polar body extrusion (Jo et al., 2015). It was shown that DNA in proximity to the cortex is sufficient to induce actin cap formation (Deng et al., 2007). Since the spindle with the DNA was intact and correctly located to the cortex in PM oocytes (**Fig. 3c**), we next analyzed the actin cap formation. The control and PM oocytes were stained with Phalloidin, a probe which visualizes F-actin (**Suppl. Fig. 4b** and **Fig. 4b**). We detected actin at the oocyte periphery in control meta- and anaphase oocytes. We also found a thick and large actin cap at the cortex overlying the chromosomes in anaphase oocytes. The actin cap in these discarded control oocytes seems to be slightly enlarged, which could be caused during mounting of the slides due to squishing of the oocytes or cellular defects in these oocytes. For example, larger abnormal actin caps are observed in oocytes that overexpressed actin (Jo et al., 2015). In PM oocytes, we also detected actin at the periphery of the oocytes, however there was no thickening of the cortical actin at the cortex in close proximity to the DNA (**Fig. 4b**). Also, no polar body protrusion was visible in the PM oocytes. In addition, F-actin was detected in bundles (**Fig. 4c**, linear fluorescence intensity analysis) throughout the whole oocytes with no specific actin layer overlapping the separating chromosomes. Increase bundle formation and lack of an actin cap was previously observed in cells lacking ARP2/3 (actin-related protein 2/3) complex, which is a component of actin cytoskeleton that initiates branching of the actin filaments (Kiehart and Franke, 2002; Nikalayevich et al., 2024; Sun et al., 2011; Wang et al., 2014). In summary, these results reveal that formation of the actin cap was faulty and actin cytoskeleton structures were dysregulated in PM oocytes. Deficiency in actin cap formation inhibits polar body formation explaining the absence of polar body protrusion in PM oocytes.

**Figure 4.**
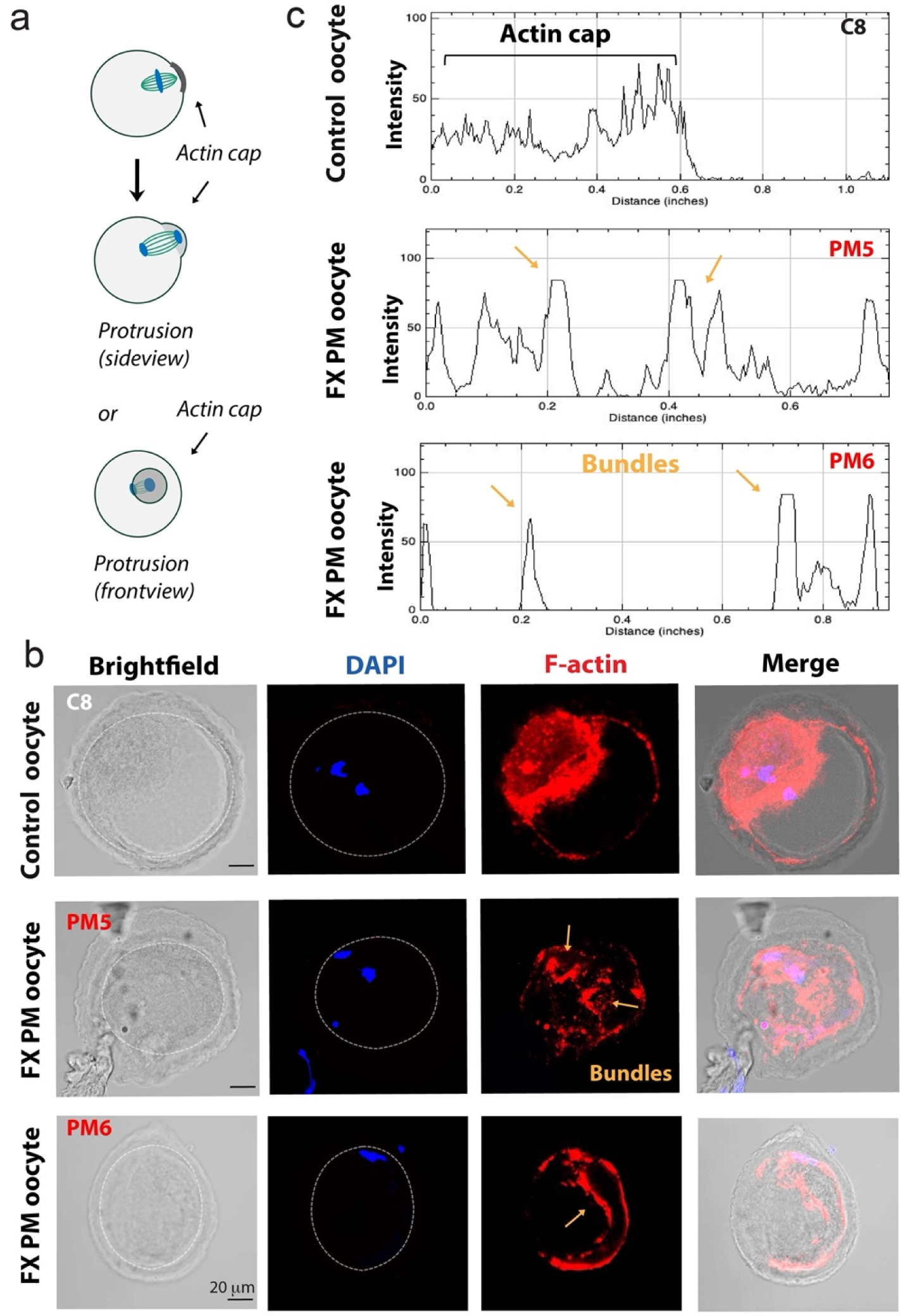
Lack of actin cap and increased bundle actin filament formation in PM oocytes. **a**) Schematic representation of actin cap formation. **b)** F-actin staining with Phalloidin (red), a high affinity F-actin probe. The DNA is visualized by DAPI (blue) staining in control and two PM oocytes. **c)** Linear fluorescence intensity analysis was used to quantify the Intensity of the immunofluorescence signals of the F-actin staining. Yellow arrows indicate the bundle actin formations.

### Elevated Profilin1 and FMRP levels result in dysregulation of the actin cytoskeleton and polar body protrusion defects in PM oocytes

In mammalian oocytes various kind of actin-based cytoskeleton structures are formed, such as bundled and branched actin filaments. These actin filament structures have specific cellular functions. One of the factors that regulate the formation of these structures is profilin1 (Davidson and Wood, 2016; Rotty et al., 2015; Suarez et al., 2015). Profilin1 is an actin binding protein which catalyzes the exchange of ADP to ATP in actin and accelerates actin polymerization. It was reported that profilin1 restrains Arp2/3 complex-mediated actin polymerization (Suarez et al., 2015). Arp2/3 induces formation of branched actin filaments that are needed for membrane protrusions (Rotty et al., 2015; Suarez et al., 2015). In addition, the profilin1 mRNA was found to be a target of FMRP (Reeve et al., 2005; Rockwell and Hongay, 2019). To test whether altered profilin1 and FMRP concentration causes lack of protrusions in PM oocytes we stained control and PM oocytes for profilin1 and FMRP (**Fig. 5a**). All analyzed oocytes were retrieved from women that were 31 years old. In control human oocytes we detected profilin1 at the cortex. In contrast in PM oocytes, we detected profilin1 not only at the cortex but throughout the oocytes (**Fig. 5a-b**). In addition, we found that profilin1 was significantly increased in PM oocytes (**Fig. 5c**). These results explain the increase formation of bundle actin filaments and the defects in membrane protrusion at the cortex in the PM oocytes.

**Figure 5.**
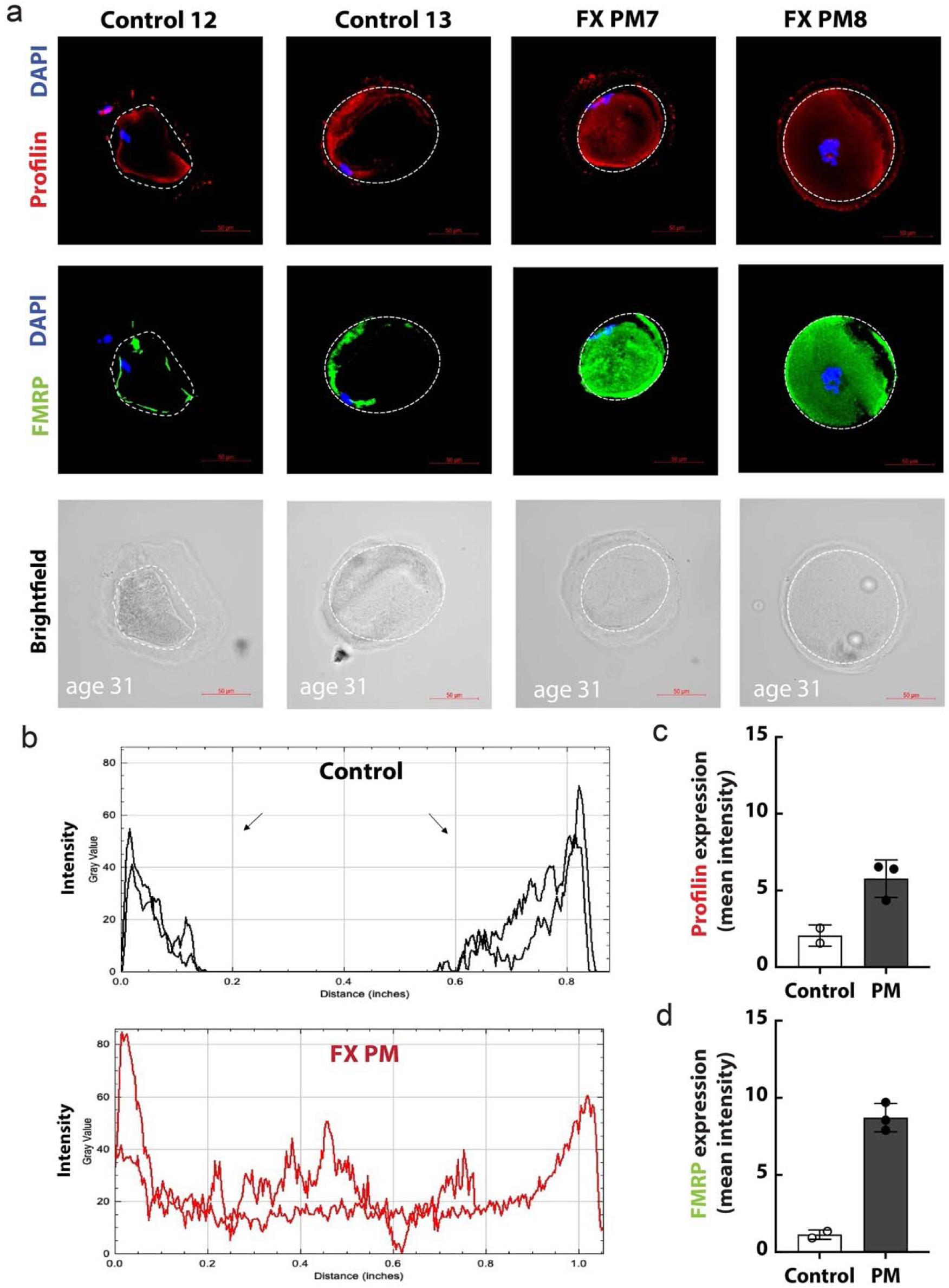
Increase in Profilin1, caused by elevated FMRP level, leads to protrusion defects in PM oocytes. **a)** Control and PM oocytes are stained with FMRP antibody (green) and profilin1 antibody (red). DNA was visualized with DAPI (blue). **b)** Linear fluorescence intensity analysis was used to quantify the Intensity and location of the profilin1 staining (from oocytes above shown in **a**) in control and PM oocytes. **c-d)** Mean intensity of profilin1 (**c**) and FMRP (**d**) staining are shown. Standard deviations are indicated. All analyzed oocytes were retrieved from women that were 31 years old.

Next, we were wondering if a reduction in FMRP expression, which was observed in PM cells (Kenneson et al., 2001) and could be responsible for the elevated profilin1 level in PM oocytes. FMRP is a known negative regulator of translation and is expressed in fetal human ovaries and in mice oocytes (Hoffman et al., 2012; Sherman et al., 2014). It was shown that in drosophila and mouse FMRP binds directly profilin1 mRNA and regulates the profilin translation as well as the actin cytoskeleton cellular architecture (Michaelsen-Preusse et al., 2016; Reeve et al., 2005). Surprisingly, we found no reduction but a significant increase in FMRP expression in PM oocytes (**Fig. 5a** and **5d**). These results reveal that FMRP is expressed in human oocytes and the FMRP translation is not constrained in PM oocytes as seen in other PM cells. The elevated FMRP expression is probably due to the increased amount of FMR1 mRNA in PM cells (Man et al., 2017; Tassone et al., 2007). FMRP was located throughout the ooplasm in PM oocytes. In contrast, in control oocytes FMRP was mostly detected at the cortex like profilin1 (**Suppl. Fig. 5a-b**). Recently, it was discovered that FMRP can also stimulates the mRNA translation (Greenblatt and Spradling, 2018). In both, control and PM oocytes, majority of FMRP colocalized with profilin1. Altogether, these findings propose that due to an increase in FMRP expression the profilin1 level is elevated, which leads to dysregulation of actin cytoskeleton structures and the protrusion defects in PM oocytes (**Fig. 7**). The failure to extrude the first polar body causes a reduction in mature MII oocytes and FXDOR/FXPOI in PM carriers.

### Dysregulation of Profilin1 and FMRP leads to actin cytoskeleton defects in aging oocytes

Fertility and reproductive aging are linked to a decline in the quantity and quality of oocytes leading to a diminished ovarian reserve with advanced maternal age (35-40) (Telfer et al., 2023). Recently discoveries were made explaining the decrease in oocytes quality and the increase in oocytes with aneuploidy. However, the reason for the decline in number of mature oocytes with advance age it is still not clearly understood. Since cytoskeleton structure are important for oocyte maturation and polar body protrusions, we asked whether dysregulation of the cytoskeleton proteins and profilin1 could prompt a decline of mature oocytes in women with advanced age. Therefore, we analyzed human oocytes from patients of age 37 and compared to patients of age 31. First, we analyzed the profilin1 and FMRP level in these oocytes. As observed before, profilin1 and FMRP were located predominantly at the cortex in oocytes from women age 31 (**Fig. 6a** and **Fig. 5**, control 12 and 13). However, we found an upsurge in profilin1 protein that expanded from the cortex into the oocyte cytoplasm in aged oocytes (age 37, **Fig. 6a-b, Suppl. 6a**). Most FMRP and profilin1 signal overlapped or were in close proximity in these human oocytes (**Suppl. Fig. 6b-c**). Also as expected, since these oocytes had normal repeat size, there was no major increase in FMRP expression (**Fig. 6c**). Nevertheless, we did observe an alteration in the FMRP location in aged oocytes. We detected FMRP not only at the cortex but also in foci in the oocyte cytoplasm (**Fig. 6a** and **d**). These findings suggest that FMRP potentially activates profilin1 translation at the cortex and in the ooplasm, which leads to elevated profilin1 expression in the cytoplasm of aged oocytes.

**Figure 6.**
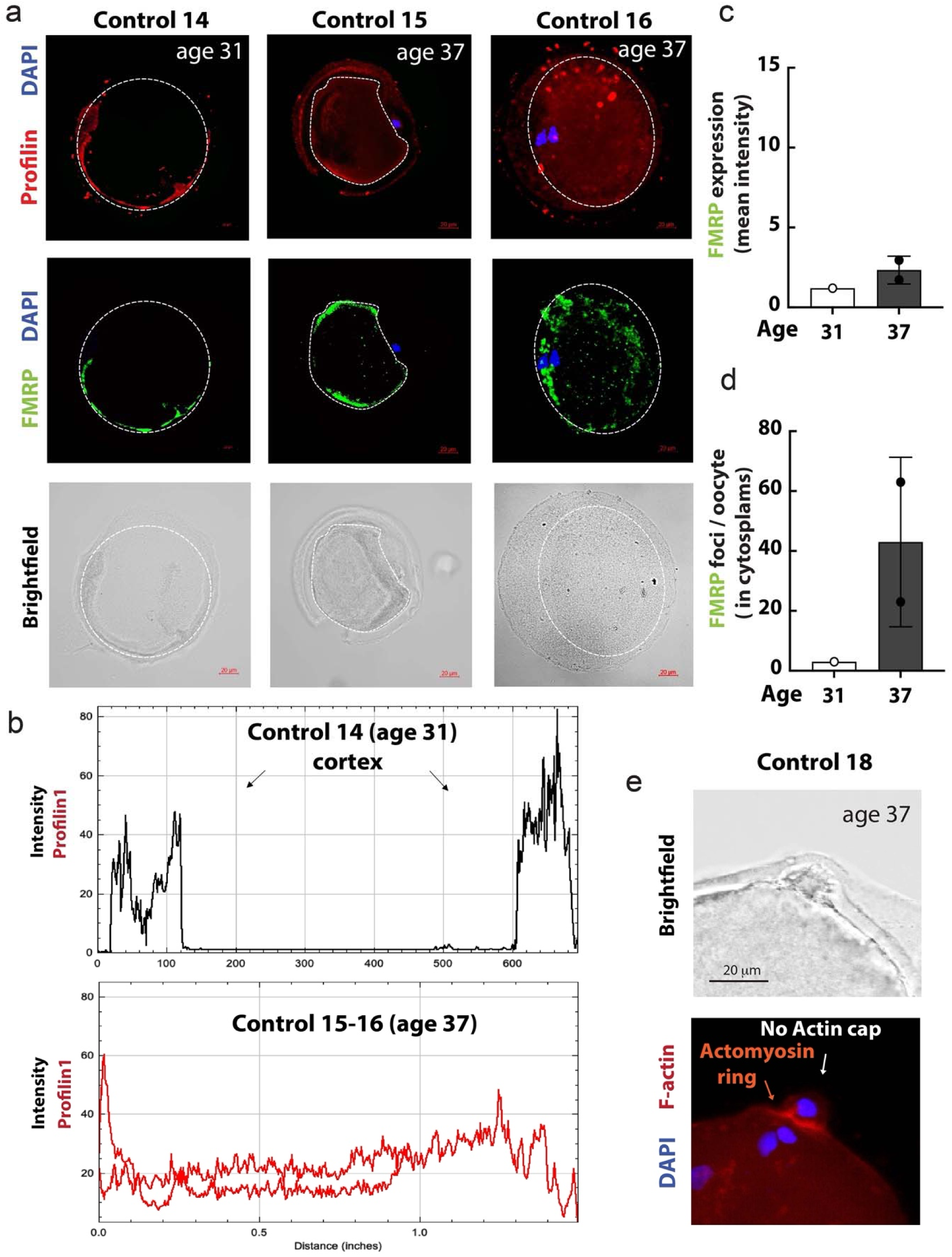
Increase in Profilin1, caused by FMRP foci, leads to actin cytoskeleton defects in aged oocytes. **a)** Control young (31 years old) and aged (37 years old) oocytes are stained with FMRP antibody (green) and profilin1 antibody (red). DNA was visualized with DAPI (blue). **b)** Linear fluorescence intensity analysis was used to quantify the Intensity and location of the profilin1 staining (from oocytes above shown in **a**) in control and PM oocytes. **c-d)** Mean intensity of FMRP (**c**) and FMRP foci in oocyte cytoplasm (**d**) are quantified. Standard deviations are indicated. **e)** F-actin staining with Phalloidin (red) and the chromosomes are visualized by DAPI (blue) staining. No actin cap was detected but the contractile (actomyosin) ring is formed.

Next, we analyzed cell cycle phase (localization of the chromosomes) and if protrusion of the first polar body occurred in aged oocytes. DAPI staining showed that all oocytes were in anaphase I. We found that two oocytes from four (age 37) showed no protrusion (**Suppl. Fig. 6d**, control 16 and 17). However, the other two oocytes from women with advanced maternal age (**Suppl. Fig. 6d**, control 15 and 18) had a projection at the cortex that contained chromosomal DNA. We stained one of these oocytes for F-actin probe Phalloidin to analyze if the actin cap is formed and if the projection is a polar body protrusion (**Fig. 6e**). Surprisingly, although no actin cap was formed, aged oocytes contained a contractile ring. It is known that the contractile ring is composed of myosin and actin bundles. This actomyosin ring was formed perpendicular to the spindle axis and formation occurred after chromosome separation (**Fig. 6e**). These results reveal that the spindle and actin bundle structure formation is not inhibited in oocytes with advanced maternal age (**Fig. 7**). However, because of dysregulation of the actin cytoskeleton, branched actin structures and actin cap formation are defective, which constrains the establishment of a proper polar body in oocytes with advanced maternal age.

**Figure 7.**
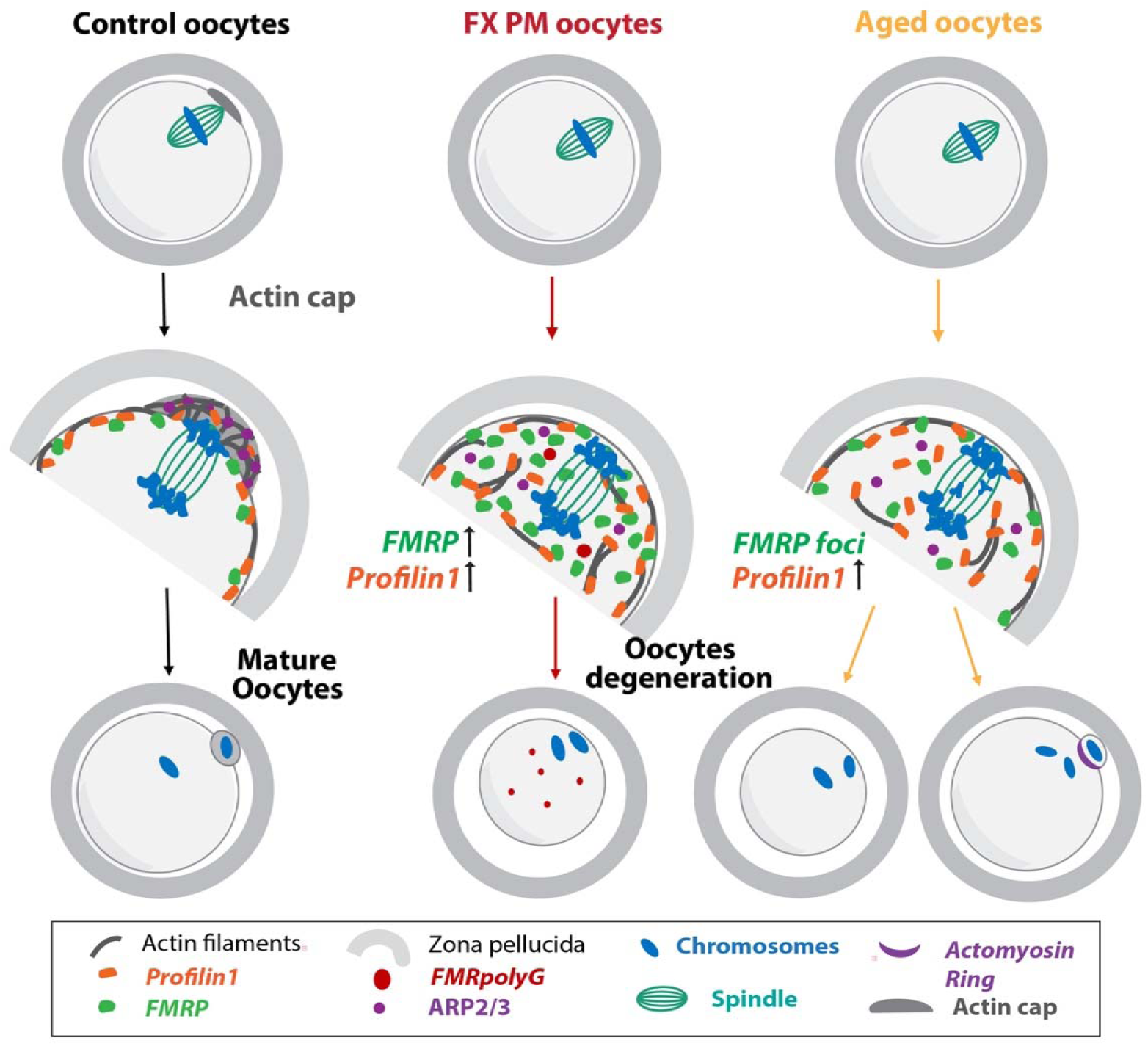
Model showing that dysregulation of FMRP and profilin1 cause defects during anaphase I that inhibit oocyte maturation in PM carriers and aged oocytes. After germinal vesicle breakdown, the spindle forms and actin filaments are dispensed evenly at the oocyte cortex. In metaphase I the spindle moves to the oocyte cortex. After spindle reaches the cortex actin accumulates to form an actin cap that overlays the spindle. In anaphase I chromosomes pair separate, and one set of chromosomes is pulled into the protruding polar body. FX PM oocytes reach anaphase I and the chromosomes are separated, however due to defects in actin cap formation no polar body is extruded and the oocytes fail to mature. Actin cap formation is constrained due to an increase in profilin1 that is upregulated by elevated FMRP level in PM oocytes. These findings propose that due to an increase in FMRP in PM oocytes, the formation of actin cytoskeleton structures is dysregulated causing polar body protrusion defects. Similar oocytes in women with advance maternal age also show an increase in profilin1, FMRP foci and actin cytoskeleton defects. The failure to extrude the first polar body leads to reduction in mature oocytes and FXDOR/FXPOI in PM carriers and in women with advanced maternal age.

## Discussion

The mechanisms and defects causing diminished ovarian reserve and ovarian dysfunction in PM carriers and women with advanced maternal age are unknown. Elucidating these mechanisms responsible for diminished ovarian reserve have proven to be extremely difficult, largely due to the scarcity of human oocytes and ovarian tissue. Due the detrimental effect on their reproductive potential, more PM carriers and women with advanced maternal age choose to undergo assisted reproductive technology (ART) to conceive a healthy child. While it is known that PM carriers may be at increased risk for lower oocyte yield, less is known about the mechanisms causing oocyte development defects in these carriers. Thus, most knowledge about the probable causes of FXDOR/FXPOI has been derived from animal model systems (Sherman et al., 2014). Studies from these model systems propose that molecular abnormalities within the ovaries or the establishment of the primordial follicle pool could be disturbed (Pepling, 2006). However, in a murine model, it was found that PM mice had the same initial primordial pool as unaffected mice, but a rapid decline of follicles of all classes occurred (Hoffman et al., 2012). It was also observed that PM mice had oocytes with increased FMR1 mRNA expression and elevated ubiquitin level. Compared to unaffected mice, PM mice also showed evidence of ovarian atrophy and damage. In humans, in a retrospective study, it was found that PM carriers had a lower number of blastocysts (Hutchinson et al., 2018). We studied human oocytes and focused our analysis on PM patients that carry 55 to 90 CGG repeats, because PM carriers with 70-90 CGG repeats have the lowest ovarian reserve (Lekovich et al., 2018; Man et al., 2017). We found reduction in both the number and the proportion of MII oocytes in PM carriers with mid-range repeat size, but no major differences in the number or proportion of GV and MI oocytes with increasing repeat length. Our results suggest that similar to studies in mice the primordial follicle pool is not affected but the oocyte maturation is constrained leading to reduced number of mature MII oocytes in PM carriers.

It was suggested that molecular abnormalities within the ovary or granulosa cells may contribute to FXDOR/FXPOI. One study found increased FMR1 mRNA expression in human granulosa cells, which were isolated after oocyte retrieval (Friedman-Gohas et al., 2020). Another study found increased translation and production of toxic FMRpolyG protein inclusions in ovarian stroma cells in one PM carrier (Buijsen et al., 2016), which could cause ovarian damage and FXPOI. While these reports offer insight into defects found in PM carriers, there is no study evaluating human PM oocytes. We analyzed human oocytes and detected cellular abnormalities, including FMRpolyG protein inclusions, irregular ubiquitin staining and DNA breaks (**Fig. 2**). Surprisingly, our results reveal that despite these cellular abnormalities, PM oocytes enter anaphase I and are able to pass the SAC and spindle positioning checkpoint, however they are unable to complete meiosis I (**Fig. 3**). These findings are consistent with our observation that there is no difference in the number of MI oocytes, but there is a reduced number of MII oocytes in PM patients (**Fig. 1** and **Suppl. Fig.1**). Furthermore, in anaphase I we found that the protrusion of the polar bodies was not initiated in PM oocytes, despite the spindle being at the cortex in pole-first direction. The separation of the homologue chromosome pairs, however, was not affected. It was shown that DNA proximal to the cortex is sufficient to induce actin cap formation (Deng et al., 2007). Conversely, we observed that PM oocytes fail to form a proper actin cap (**Fig. 4**) (Dehapiot et al., 2013) due to dysregulation of the actin cytoskeleton which explains the decline in mature MII oocytes and FXDOR/FXPOI.

Accurate coordination of the assembly/disassembly of the actin cytoskeleton is a crucial requirement to shape cells and perform specific cell functions. Actin cap and branched actin structures are needed for protrusions. Profilin1 and other regulatory proteins are essential for organization of the actin cytoskeleton. It was shown that the ratio of profilin1 regulates the formation of actin filament structures (Davidson and Wood, 2016). Profilin1 mRNA is a target of FMRP (Reeve et al., 2005; Rockwell and Hongay, 2019). FMRP is expressed in rat and mice oocytes as well as in fetal human ovaries (Hoffman et al., 2012; Sherman et al., 2014). Surprisingly, we found an increase in FMRP in PM oocytes (**Fig. 5**) and not a reduction how it was observed in mouse oocytes (Hoffman et al., 2012). These results suggest that FMRP translation is not inhibited in human PM oocytes. We also found that FMRP is located predominately at the cortex in human cells, which is different to mice but similar to bovine oocytes (Karen Nenonene et al., 2023), suggesting that FMRP has distinct functions in oocytes which differs between mammalian species. In addition, we found an increase in profilin1 level, which due to a strong colocalization with FMRP suggest that FMRP stimulates the translation of profilin1 mRNA in human oocytes. This is also supported by reduced proflin1 level, in a mouse model that lacks FMRP (Michaelsen-Preusse et al., 2016). Initial research showed that FMRP suppresses the translation of mRNAs into proteins (Laggerbauer et al., 2001; Li et al., 2001). However, recently it was found that FMRP also activates the translation of mRNAs in drosophila oocytes, particularly of large mRNAs (Greenblatt and Spradling, 2018). Two other studies also confirmed this finding (Flanagan et al., 2022; Seo et al., 2022) that FMRP activates the translation of mRNAs and that this novel function of FMRP, which was also observed both in *Drosophila* and mouse, is evolutionary conserved (Flanagan et al., 2022).

It is known that the quality (such as an increase in aneuploidy (Charalambous et al., 2023)) and quantity of oocytes decreases with advanced maternal age. However, the mechanisms causing the deterioration of mature oocytes it is not clearly understood (Telfer et al., 2023). So far, it is assumed that follicular atresia and a declining oocyte pool causes the reduced number of oocytes with advanced maternal age. However, at age 35-39 when the fertility starts to decline rapidly there are still enough oocytes left in the pool (at age 37 around 25,000 oocytes left according to ACOG). We found that maturation of the oocytes in women with advanced maternal age (37 years) is hindered by deficiencies in forming branched actin cytoskeleton structures needed for accurate extrusion of the first polar body. The cause is similar to the mechanism in PM carriers. Aged oocytes had an increase in profilin1 and FMRP foci that expanded from the cortex into the oocyte cytoplasm leading to dysregulation of the cytoskeleton and lack of actin cap formation (**Fig. 6**). Interesting, the migration and formation of the spindle and bundle actin structure such as found in actomyosin ring was not affected. These results confirm that cytoskeleton formation is dysregulated and that specifically the branched actin structure formation is defective.

It remains to be determined whether other PM tissues and PM cells have also an increase in FMRP instead a decrease in FMRP. Also, dysregulation of FMRP and actin cytoskeleton abnormalities might play a role in the disease pathology in other PM tissues. In addition, it is not clear if the observed FMR1polyG aggregates, DNA breaks and abnormal ubiquitin level contribute to the FXDOR/FXPOI pathology or are just consequences of the CGG repeat expansions. Also, it is possibility that the demise of granulosa cells and a subsequent lack of nutritions and factors, that are important for oocyte maturation, could promote degeneration of PM oocytes. In summary, this study for the first time analyzes human oocytes obtained from PM carriers and reveals the mechanism and defects leading to FXPOI/FXDOR. These results show that deficiencies during anaphase I and abnormal cytoskeleton structure formation hinder oocyte maturation leading to a reduction of mature MII oocytes and FXDOR/FXPOI symptoms in PM carriers (**Fig. 7**). Our results also show that FMRP and profilin1 play a pivotal role in human oocyte maturation and dysregulation of both proteins and the cytoskeleton is a part of the aging process leading to less mature oocyte with advance age.

## Materials and Methods

### Retrospective analysis: Inclusion and Exclusion Criteria of IVF Patients

The institutional review board at Weill Cornell Medical College approved our study protocol. For this retrospective cohort study, all PM carriers (55-200 CGG repeats) and intermediate allele carriers (45-54 CGG repeats) undergoing IVF at the Ronald O. Perelman and Claudia Cohen Center for Reproductive Medicine were assessed for potential inclusion. PM carriers with unknown number of CGG repeats, with two affected alleles or who had cancelled cycles were excluded. The control group included age-matched patients undergoing IVF for preimplantation genetic testing for monogenic disorders (PGT-M) during the same time period. The Institutional Review Board of WCM approved the current study.

### Clinical and Laboratory Protocols

Ovarian stimulation, trigger for final oocyte maturation, and oocyte retrieval were performed per our standard protocols (Huang and Rosenwaks, 2014). Ovarian stimulation was carried out to maximize follicular response while minimizing the risk of ovarian hyperstimulation syndrome (OHSS). The initial gonadotropin dose was based on age, weight, antral follicle count, AMH and previous response to stimulation. Ovarian stimulation was carried out with a combination of gonadotropins (Follistim, Merck, Kenilworth, NJ, USA; Gonal-F, EMD-Serono, Geneva, Switzerland; and/or Menopur, Ferring Pharmaceuticals Inc, Parsippany, NJ, USA), with ovulation being suppressed with once daily 0.25 mg Ganirelix Acetate (Merck, Kenilworth, NJ, USA) or Cetrotide (EMD-Serono, Geneva, Switzerland) injections (Huang and Rosenwaks, 2014).

Human chorionic gonadotropin (hCG) was used as the ovulation trigger in the majority of cycles during the early years of the study period and gonadotropin-releasing hormone (GnRH) agonist triggers were incorporated into clinical practice in the latter years of the study period. Novarel (Ferring Pharmaceuticals Inc, Parsippany, NJ, USA) or Pregnyl (Merck, Kenilworth, NJ, USA) was administered according to a sliding scale (10000 IU for Estradiol (E_2_) <1500 pg/mL, 5000 IU for E_2_ 1501–2500 pg/mL, 4000 IU for E_2_ 2501–3000 pg/mL, and 3300 IU for E_2_>3001 pg/mL) when given as the sole trigger (Huang and Rosenwaks, 2014). The trigger was given when the two lead follicles attained at least a mean diameter >17 mm. Oocyte retrieval was performed with transvaginal ultrasound guidance under conscious sedation approximately 35– 36 hours after hCG administration. All retrieved oocytes were exposed to 40 IU recombinant hyaluronidase (Cumulase, Halozyme Therapeutics Inc., San Diego, CA, USA) to remove the cumulus-corona complex. Measurements of serum E_2_, progesterone (P) and hCG were performed in our center’s laboratory using the IMMULITE 2000 Immunoassay System (Siemens, Berlin, Germany). The sensitivity of the E_2_, P and hCG assay is 20 pg/mL, 0.2 ng/mL, and 0.4 mIU/mL, respectively. All intra- and inter-assay variation coefficients are <10.

### Study Variables and Statistical Analyses

Demographic and baseline characteristics recorded for all recipients included age at retrieval, BMI (kg/m^2^), gravidity, parity, AMH level (ng/ml), number of CGG repeats, and number of previous IVF cycles. The primary outcome was oocyte maturity after oocyte retrieval, which was evaluated by percent of metaphase II (MII) oocytes and percent of oocytes arrested in prophase I or metaphase I (MI). The degree of oocyte maturity was then compared to the number of CGG repeats. Patients were divided into three groups based on repeat number (45-54, 55-68 and 69-85 CGG repeats). This grouping was chosen due to prior data suggesting that PM carriers with 70-90 CGG repeats are at greater risk of FXPOI. Secondary outcomes included AMH level, number of oocytes retrieved, days of stimulation and total gonadotropin dosage.

Descriptive statistics were used to characterize the study sample with respect to demographic and clinical factors of interest. Continuous variables are expressed as mean (standard deviation) and categorical variables are expressed as n (%). Where appropriate, unpaired t-test was performed for statistical analysis. All *p*-values are two-sided with statistical significance evaluated at the 0.05 alpha level. All analyses were performed in Stata BE 17.0 (StataCorp LLC) and Prism (GraphPad Software year 2023).

### Immature oocyte collection and Immunostaining

We only collected discarded immature oocytes from patients going through IVF. The study was approved by the Institutional Review Board of Weill Cornell Medicine. The embryologist denuded the oocytes before our analysis and categorized the oocyte as mature (MII) or immature (GV or MI). PM oocytes were collected and fixed on positively charged slides and frozen at -80°C and stored until future use. The slides were kept on ice before use and were allowed to thaw for a few minutes only. Briefly, the oocytes were fixed with the fixation-permeabilization buffer for 30-40 min at RT (4% paraformaldehyde, 200 mM PIPES-pH 6.8, 200 mM MgCl2, 10 mM EGTA, 0.2% Triton X). The slides were washed gently with PBS and the oocytes were viewed continuously during the process under the microscope to ensure that oocytes were not disturbed or washed away. The oocytes were blocked with 1%BSA and 0.3% Triton X in PBS (PBST) for 1 h and then incubated overnight with primary antibody at 4°C. The oocytes were washed thrice with PBS and followed by incubation with secondary antibody (Alexa 488 or 594; 1:1000 in PBST) at room temperature for 1 h. After PBS wash, the slides were mounted with ProLong Gold Antifade Mountant with DAPI. Primary antibodies used in immunofluorescence studies were γ-H2AX (Phospho-Histone H2A.X Ser139-D7T2V, Mouse mAb #80312 Cell Signaling Technology; 1:1000), Ubiquitin Polyclonal Antibody (Invitrogen #PA1-26088; 1:150), anti-FMR1polyG Antibody (clone 2J7, #MABN784 Millipore, 1: 100), Prolifin 1 (PFN1 Monoclonal Antibody ThermoFisher, #67390-1-IG, 1:100), Anti-FMRP antibody (Polyclonal Antibody Abcam, #ab17722, 1: 200) and Alexa Fluor 546 phalloidin probe (# A22283 Invitrogen, 5U/ml for 20 min at RT). Pictures of the stained oocytes were taken with Zeiss confocal microscope (Model LSM710). The signal intensity of the immunostaining across the oocytes were analyzed by lines plots using ImageJ.

## Acknowledgment

This work was supported by R35GM152228 (J.G.)

---

**Suppl. Table 1.** Demographic and baseline characteristics of IVF patients used in this study, related to Table 1. The mean age, BMI, ethnicity, gravidity, parity, AMH and number of previous IVF cycle are displayed for control, intermediate and PM carriers. The groups are similar for gravidity, parity, and BMI. The *p*-value are shown.

| Suppl Table 1. Patient characteristics and baseline demographics |  |  |  |  |  |  |  |  |  |
| --- | --- | --- | --- | --- | --- | --- | --- | --- | --- |
|  | Premutation Group (n=34)<sup>†</sup> | Control Group (n=207)<sup>†</sup> | P-value* | Premutation Group (n=34)<sup>†</sup> | Intermediate Group (n=44)<sup>†</sup> | P-value* | Intermediate Group (n=44)<sup>†</sup> | Control Group (n=207)<sup>†</sup> | P-value* |
| Patient age | 33.8 ± 3.9 | 33.8 ± 4.0 | 0.9 | 33.8 ± 3.9 | 36.8 ± 3.7 | <0.001 | 36.8 ± 3.7 | 33.8 ± 4.0 | <0.001 |
| BMI | 20.8 ± 7.7 | 22.1 ± 8.0 | 0.2 | 20.8 ± 7.7 | 23.7 ± 4.0 | 0.003 | 23.7 ± 4.0 | 22.1 ± 8.0 | 0.01 |
| Ethnicity |  |  |  |  |  |  |  |  |  |
| Caucasian | 19 (57.6%) | 134 (67.7%) |  | 19 (57.6%) | 36 (81.8%) |  | 36 (81.8%) | 134 (67.7%) |  |
| Black | 2 (6.1%) | 24 (12.1%) |  | 2 (6.1%) | 0 (0%) |  | 0 (0%) | 24 (12.1%) |  |
| Asian | 2 (6.1%) | 7 (3.5%) |  | 2 (6.1%) | 2 (4.5%) |  | 2 (4.5%) | 7 (3.5%) |  |
| Other | 9 (27.3%) | 13 (6.6%) |  | 9 (27.3%) | 1 (2.3%) |  | 1 (2.3%) | 13 (6.6%) |  |
| Declined | 1 (3.0%) | 20 (10.1%) |  | 1 (3.0%) | 5 (11.4%) |  | 5 (11.4%) | 20 (10.1%) |  |
| Gravidity | 1.1 ± 1.6 | 0.9 ± 1.3 | 0.3 | 1.1 ± 1.6 | 1.0 ± 1.5 | 0.6 | 1.0 ± 1.5 | 0.9 ± 1.3 | 0.5 |
| Parity | 0.3 ± 0.6 | 0.4 ± 0.7 | 0.5 | 0.3 ± 0.6 | 0.2 ± 0.5 | 0.1 | 0.2 ± 0.5 | 0.4 ± 0.7 | 0.01 |
| AMH (ng/ml) | 2.3 ± 2.3 | 3.2 ± 3.2 | 0.02 | 2.3 ± 2.3 | 3.4 ± 2.9 | 0.02 | 3.4 ± 2.9 | 3.2 ± 3.2 | 0.4 |
| Previous IVF cycle | 2.3 ± 2.9 | 1.3 ± 1.6 | 0.002 | 2.3 ± 2.9 | 1.8 ± 2.8 | 0.2 | 1.8 ± 2.8 | 1.3 ± 1.6 | 0.1 |
| <sup>†</sup> n refers to number of individual patients<br>* *p* < 0.05 is statistically significant |  |  |  |  |  |  |  |  |  |

**Suppl. Table 2.** IVF cycle outcomes, related to Table 1. The peak estradiol level, total gonadotropin dosage and days of stimulation, number of retrieved oocytes, percent of MII oocytes, GV oocytes and MI oocytes are displayed for control, intermediate and PM carriers. The *p*-value are shown.

| Suppl. Table 2. IVF Cycle Outcomes |  |  |  |  |  |  |  |  |  |
| --- | --- | --- | --- | --- | --- | --- | --- | --- | --- |
|  | Premutation Group (n=83) † | Control Group (n=249) † | p-value* | Premutation Group (n=83) † | Intermediate Group (n=96) † | p-value* | Intermediate Group (n=96) † | Control Group (n=249) † | p-value* |
| Peak E2 Level | 2235.6 ± 1625.2 | 2232.8 ± 1144.6 | 0.9 | 2235.6 ± 1625.2 | 2175.8 ± 1125.5 | 0.8 | 2175.8 ± 1125.5 | 2232.9 ± 1144.8 | 0.4 |
| Total Gonadotropin dosage (IU) | 3401.2 ± 1642.0 | 3322.6 ± 1638.6 | 0.7 | 3401.2 ± 1642.0 | 3106.8 ± 1351.6 | 0.2 | 3106.8 ± 1351.6 | 3322.6 ± 1638.6 | 0.4 |
| Days of Stimulation | 11.4 ± 2.5 | 10.8 ± 3.2 | 0.1 | 11.4 ± 2.5 | 10.0 ± 2.5 | **<0.01** | 10.0 ± 2.5 | 10.8 ± 3.2 | 0.07 |
| Number of Oocytes Retrieved | 13.1 ± 9.4 | 15.9 ± 9.5 | **0.02** | 13.0 ± 9.4 | 14.0 ± 9.6 | 0.5 | 14.0 ± 9.6 | 15.9 ± 9.7 | 0.1 |
| Number of MII Oocytes | 10.4 ± 7.8 | 12.3 ± 7.8 | **0.05** | 10.4 ± 7.8 | 10.5 ± 7.0 | 0.9 | 10.5 ± 7.0 | 12.3 ± 7.8 | **0.04** |
| Percent of MII Oocytes out of retrieved oocytes | 77.1% | 78.4% | 0.64 | 77.1% | 77.9% | 0.8 | 77.9% | 78.4% | 0.7 |
| Percent Germinal Vesicles out of immature oocytes | 55.0% | 56.8% | 0.8 | 55.0% | 39.2% | **0.02** | 39.2 % | 56.8% | **0.001** |
| Percent of MI oocytes out of immature oocytes | 32.5% | 32.5% | 0.9 | 32.5% | 43.0% | 0.2 | 43.0% | 32.5% | 0.06 |
| † n refers to number of IVF cycles<br>* p < 0.05 is statistically significant |  |  |  |  |  |  |  |  |  |

**Suppl. Fig.1.**
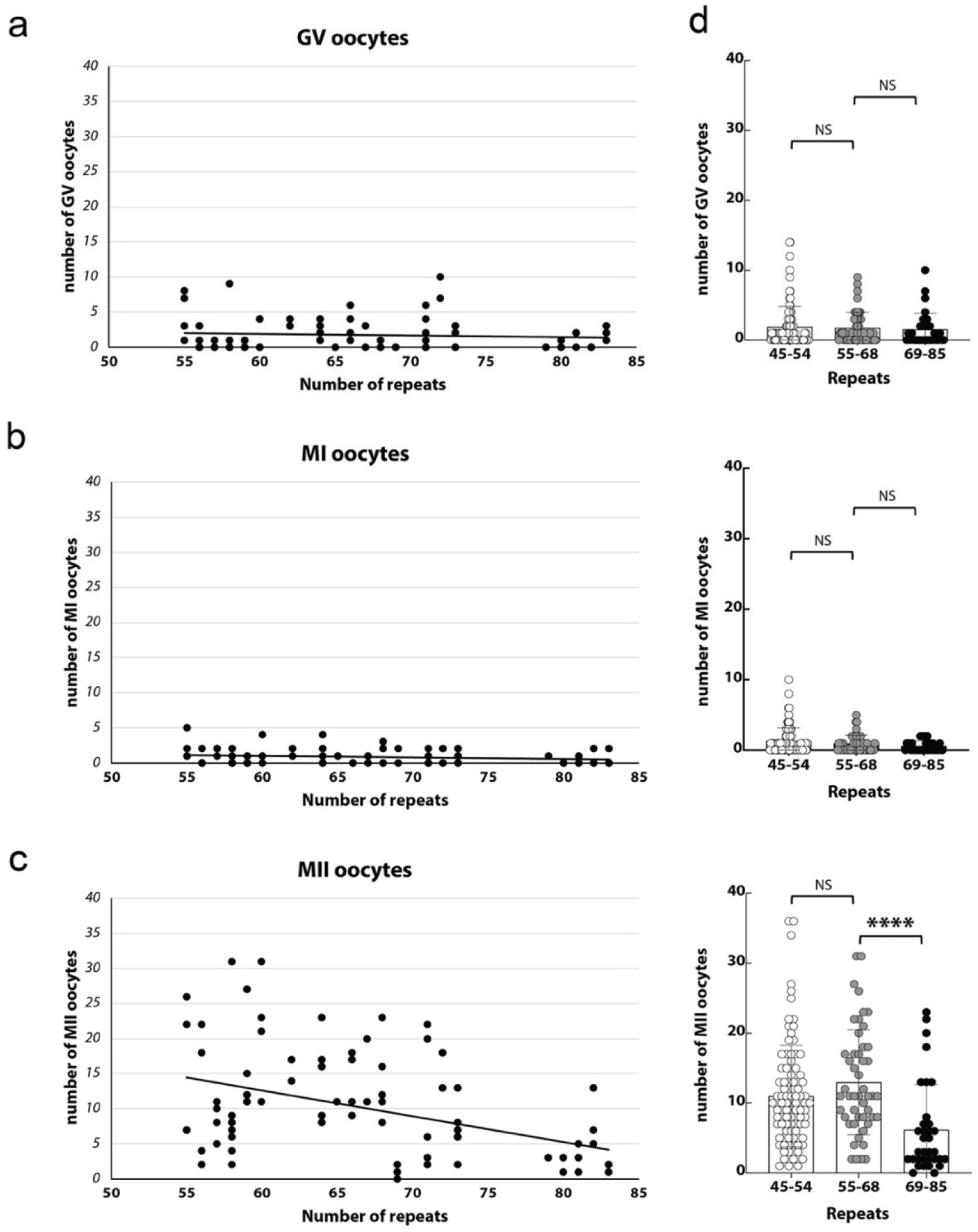
Reduction in mature MII oocytes with increasing CGG repeat length were observed in PM carriers, related to Figure 1. **a-c)** Diagrams of the total number of GV (**a**), MI (**b**) and MII (**c**) oocytes are plotted over the CGG repeat length (55 to 85 CGG repeats). **d)** PM patients were divided into three groups based on repeat number (intermediate (45-54), PM from 55-68 and from 69-85 repeats). We divided the PM carriers in two groups, low repeat number PM (55-68 CGG) and mid-range repeat number PM (69-85 CGG) group. The average of the number of GV, MI and MII oocytes are shown. Standard deviation and *p*-value (NS, not significant, * < 0.05, ** < 0.005) are indicated.

**Suppl. Fig.2.**
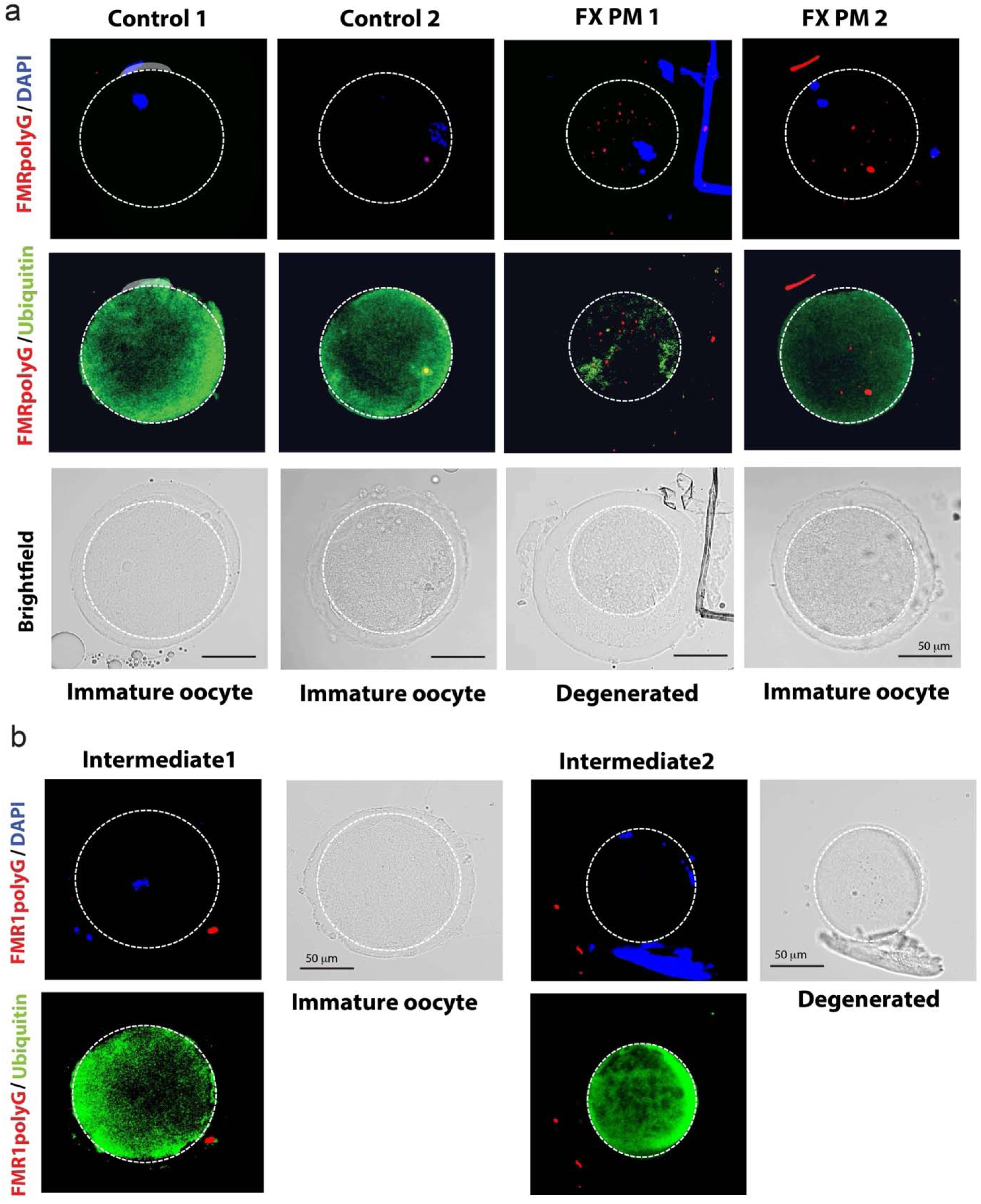
Analysis of FMRpolyG inclusions in PM and intermediate (IM) oocytes, related to Figure 2. **a)** Immature control and PM oocytes were stained with FMRpolyG antibody (red) and ubiquitin antibody (green). The DNA was visualized with DAPI. **b)** Two IM oocytes are stained with FMRpolyG antibody (red) and ubiquitin antibody (green). The DNA was visualized with DAPI.

**Suppl. Fig.3.**
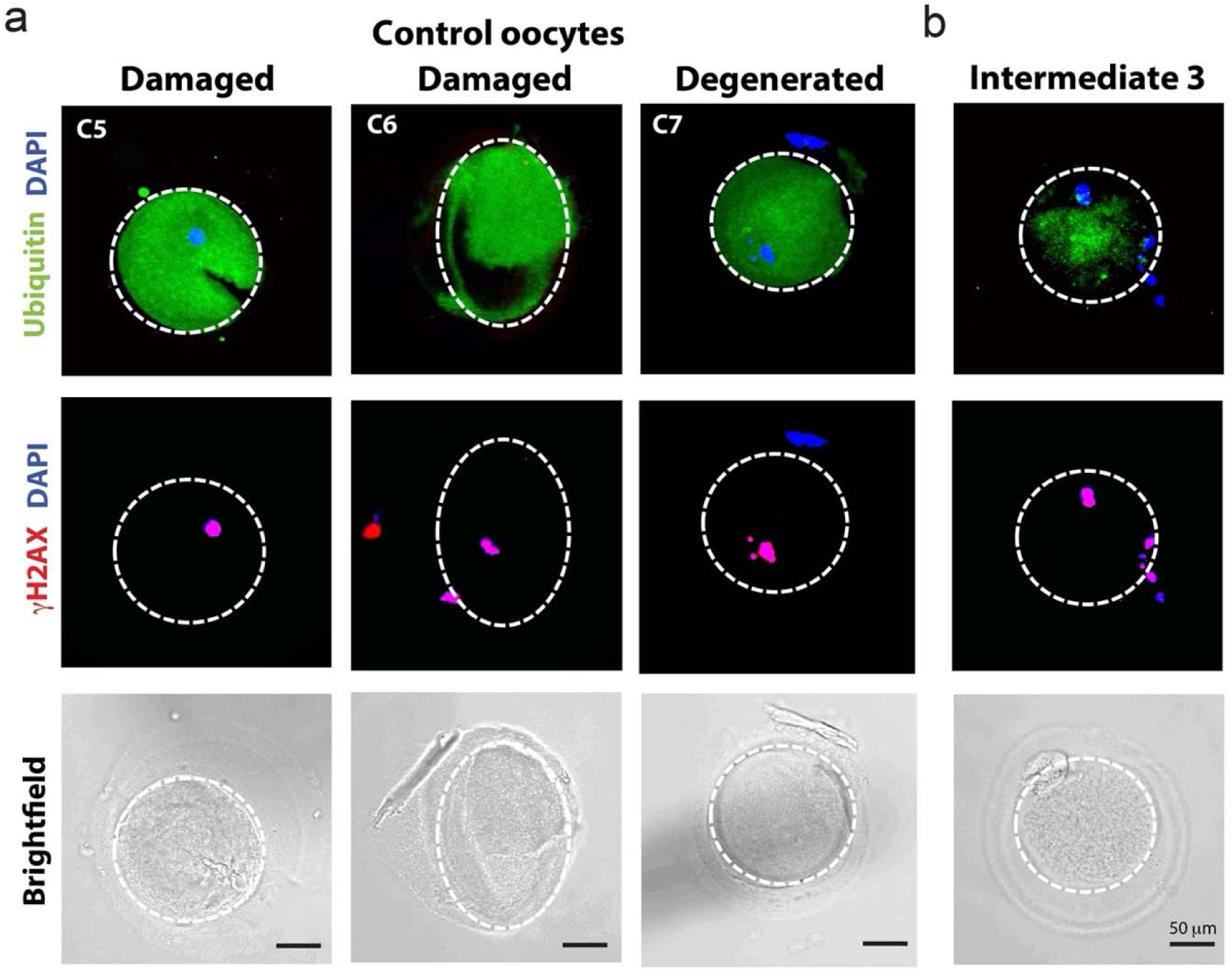
Ubiquitin staining and DNA breaks analysis in control and intermediate oocytes, related to Figure 2. **a-b).** Damaged and degenerative control (**a**) and one IM oocytes (**b**) stained with ubiquitin antibody (green), and DNA breaks with γH2AX antibody (red). DNA was visualized with DAPI (blue).

**Suppl. Fig.4.**
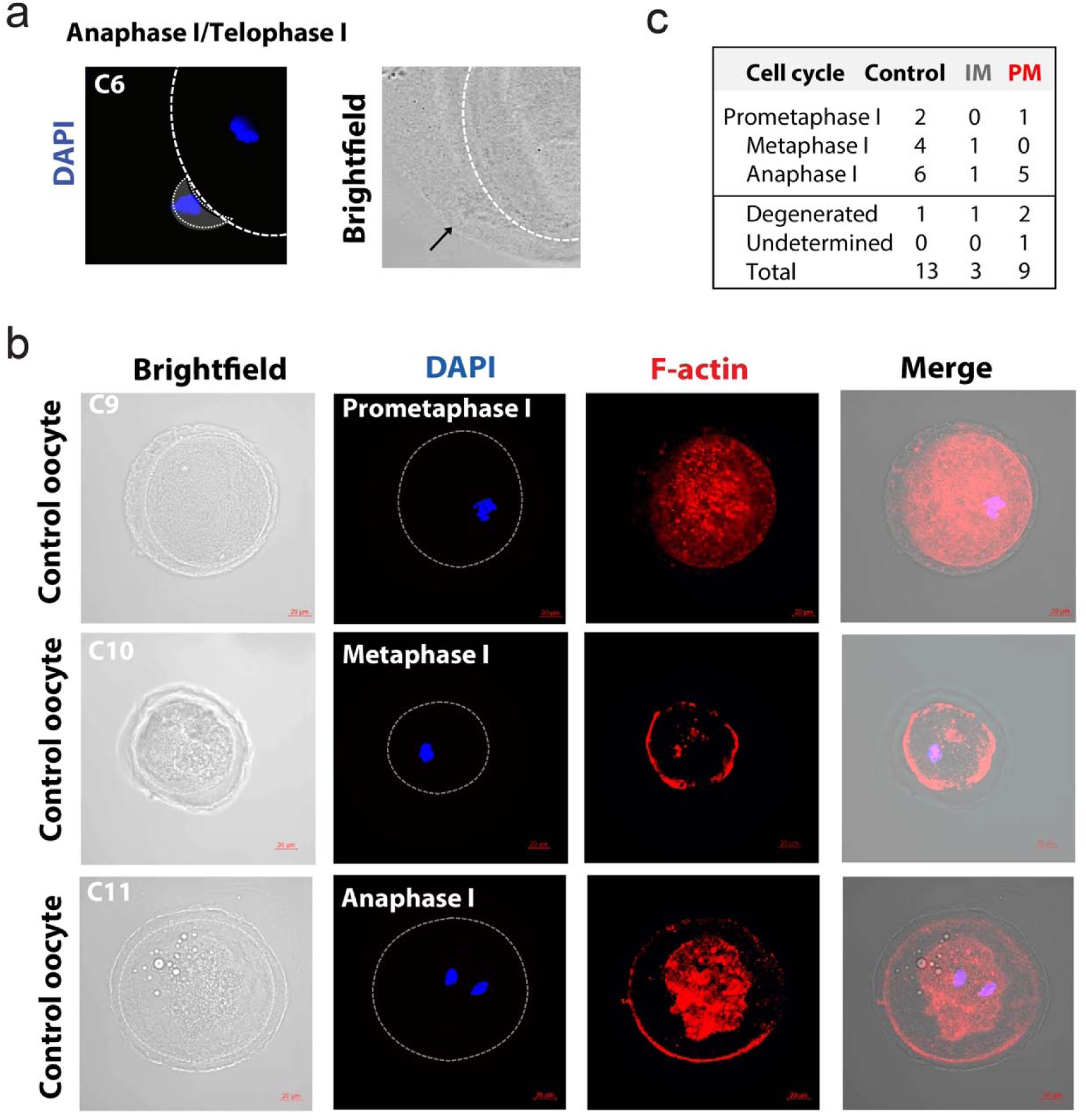
F-actin staining of control oocytes during maturation, related to Figure 3 and 4. **a)** DAPI staining and Brightfield image of a PM oocyte with visible protrusion of the first polar body (black arrow). **b)** Control oocytes in different cell cycle phases were stained with Phalloidin (red), a high affinity F-actin probe, and DAPI (blue). Images merged with brightfield are also shown. **c)** Table summarizes the total number of analyzed PM and IM oocytes in different the cell cycle phase and includes number of degenerated oocytes (and oocytes where cell cycle phase could not be determined).

**Suppl. Fig.5.**
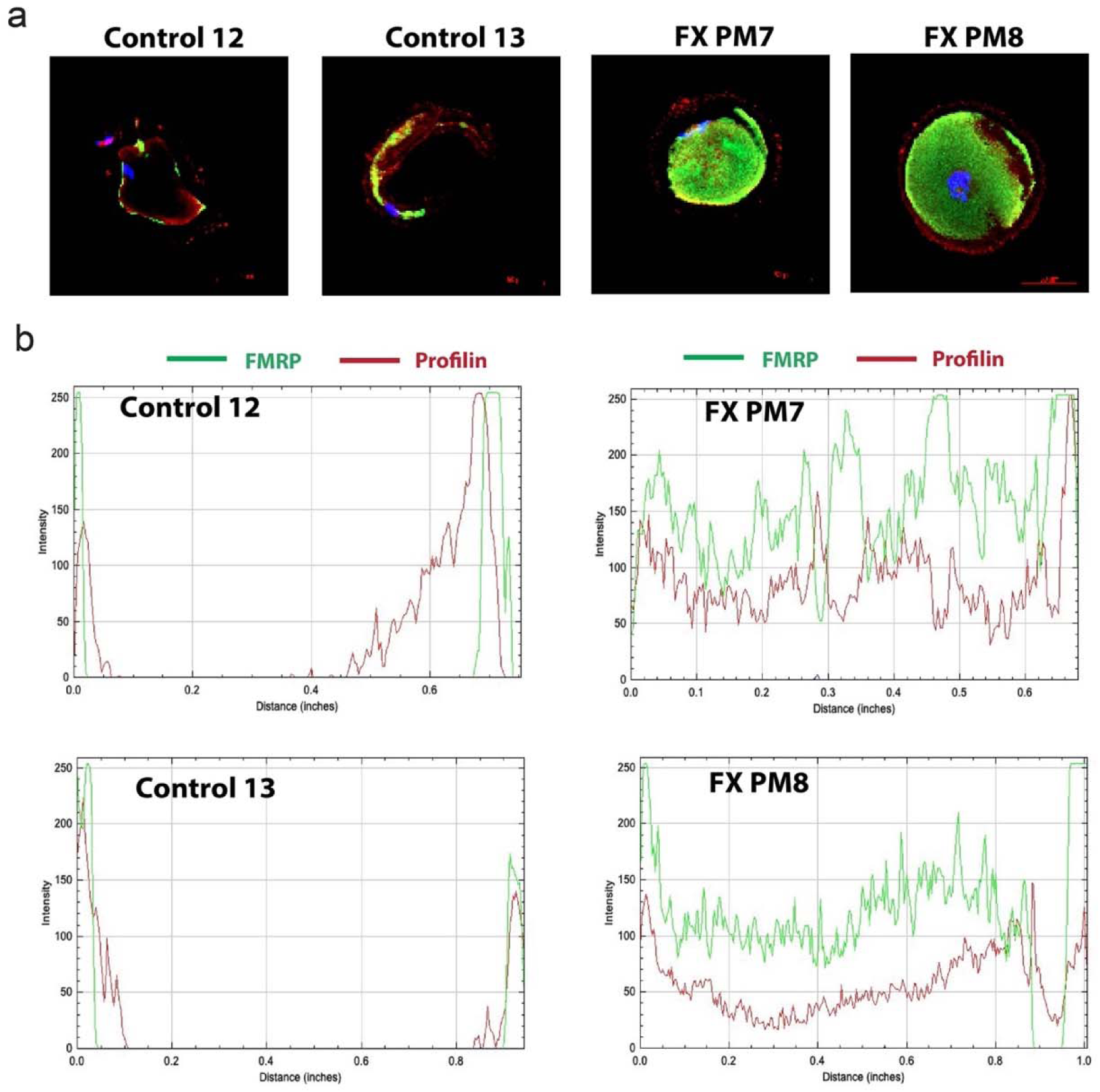
Profilin and FMRP colocalize in control and PM oocytes, related to Figure 5. **a)** Control and PM oocytes are stained with FMRP antibody (green) and profilin antibody (red). DNA was visualized with DAPI (blue). **b)** Intensity line plots of profilin staining (from oocytes above in **a**) show the location of profilin (red) and FMRP (green) in control and PM oocytes.

**Suppl. Fig.6.**
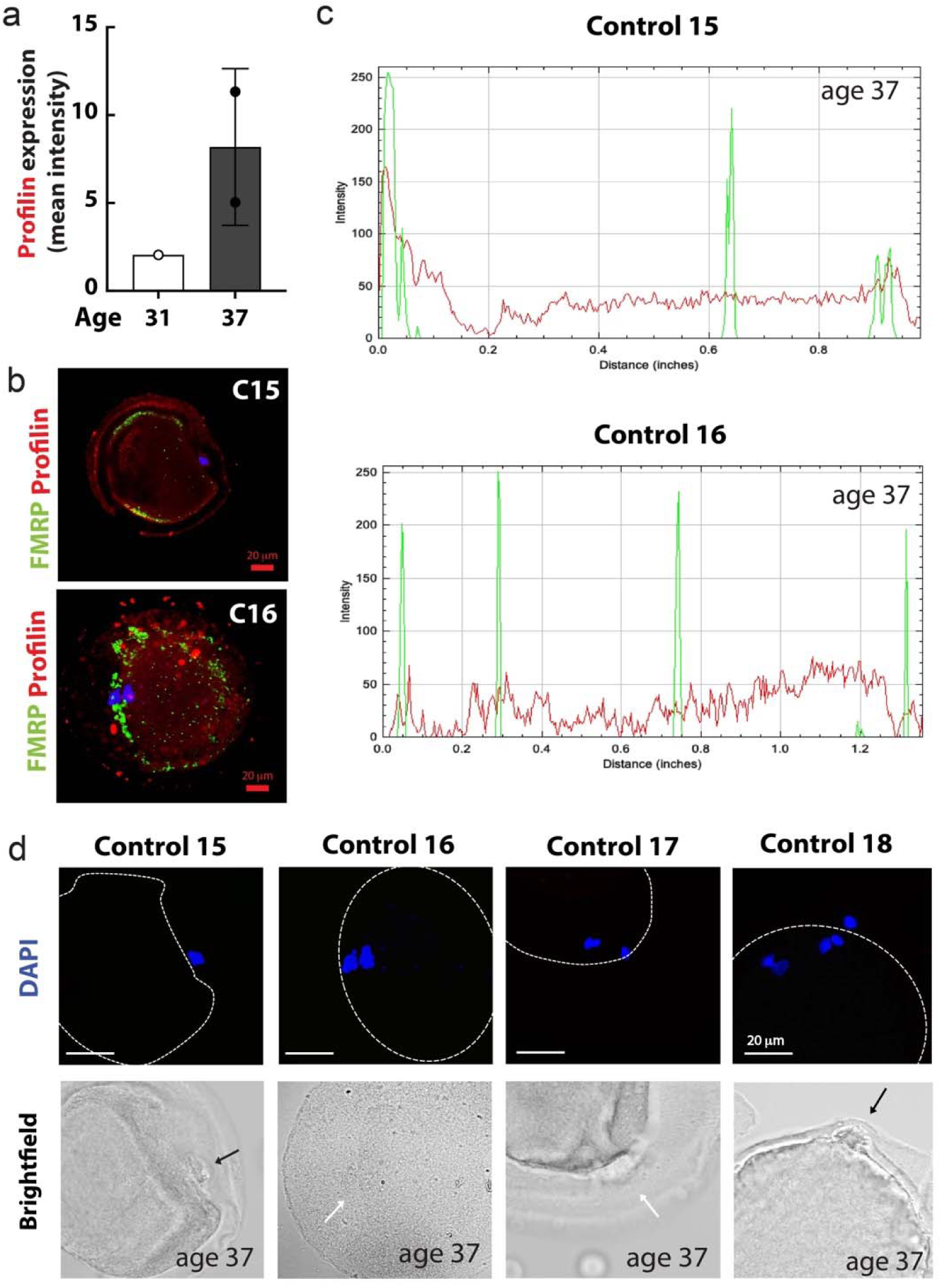
Aged oocytes were analyzed for dysregulation of proteins and protrusion defects, related to Figure 6. **a)** Mean intensity of profilin1 staining is shown (from **Fig. 6a**). Standard deviations are indicated. **b-c)** Control oocytes from women age 37 years are stained with FMRP antibody (green) and profilin1 antibody (red). DNA was visualized with DAPI (blue). **c)** Intensity line plots of profilin and FMRP staining (from oocytes in **b**) show the location of profilin (red) and FMRP (green) in control oocytes. **c)** Brightfield pictures of four control oocytes age 37, where protrusion was visible (black arrow) or absent (white arrow).

## References

Allen, E.G., Charen, K., Hipp, H.S., Shubeck, L., Amin, A., He, W., Nolin, S.L., Glicksman, A., Tortora, N., McKinnon, B., et al. (2021). Refining the risk for fragile X-associated primary ovarian insufficiency (FXPOI) by FMR1 CGG repeat size. Genet Med 23, 1648–1655.

Allen, E.G., Sullivan, A.K., Marcus, M., Small, C., Dominguez, C., Epstein, M.P., Charen, K., He, W., Taylor, K.C., and Sherman, S.L. (2007). Examination of reproductive aging milestones among women who carry the FMR1 premutation. Hum Reprod 22, 2142–2152.

Buijsen, R.A., Visser, J.A., Kramer, P., Severijnen, E.A., Gearing, M., Charlet-Berguerand, N., Sherman, S.L., Berman, R.F., Willemsen, R., and Hukema, R.K. (2016). Presence of inclusions positive for polyglycine containing protein, FMRpolyG, indicates that repeat-associated non-AUG translation plays a role in fragile X-associated primary ovarian insufficiency. Hum Reprod 31, 158–168.

Charalambous, C., Webster, A., and Schuh, M. (2023). Aneuploidy in mammalian oocytes and the impact of maternal ageing. Nat Rev Mol Cell Biol 24, 27–44.

Davidson, A.J., and Wood, W. (2016). Unravelling the Actin Cytoskeleton: A New Competitive Edge? Trends Cell Biol 26, 569–576.

Dehapiot, B., Carriere, V., Carroll, J., and Halet, G. (2013). Polarized Cdc42 activation promotes polar body protrusion and asymmetric division in mouse oocytes. Dev Biol 377, 202–212.

Deng, M., Suraneni, P., Schultz, R.M., and Li, R. (2007). The Ran GTPase mediates chromatin signaling to control cortical polarity during polar body extrusion in mouse oocytes. Dev Cell 12, 301–308.

Fabritius, A.S., Ellefson, M.L., and McNally, F.J. (2011). Nuclear and spindle positioning during oocyte meiosis. Curr Opin Cell Biol 23, 78–84.

Farrawell, N.E., McAlary, L., Lum, J.S., Chisholm, C.G., Warraich, S.T., Blair, I.P., Vine, K.L., Saunders, D.N., and Yerbury, J.J. (2020). Ubiquitin Homeostasis Is Disrupted in TDP-43 and FUS Cell Models of ALS. iScience 23, 101700.

Flanagan, K., Baradaran-Heravi, A., Yin, Q., Dao Duc, K., Spradling, A.C., and Greenblatt, E.J. (2022). FMRP-dependent production of large dosage-sensitive proteins is highly conserved. Genetics 221.

Friedman-Gohas, M., Elizur, S.E., Dratviman-Storobinsky, O., Aizer, A., Haas, J., Raanani, H., Orvieto, R., and Cohen, Y. (2020). FMRpolyG accumulates in FMR1 premutation granulosa cells. J Ovarian Res 13, 22.

Galloway, J.N., and Nelson, D.L. (2009). Evidence for RNA-mediated toxicity in the fragile X-associated tremor/ataxia syndrome. Future Neurol 4, 785.

Greaney, J., Wei, Z., and Homer, H. (2018). Regulation of chromosome segregation in oocytes and the cellular basis for female meiotic errors. Hum Reprod Update 24, 135–161.

Greenblatt, E.J., and Spradling, A.C. (2018). Fragile X mental retardation 1 gene enhances the translation of large autism-related proteins. Science 361, 709–712.

Hagerman, P.J., and Hagerman, R.J. (2004). The fragile-X premutation: a maturing perspective. American journal of human genetics 74, 805–816.

Hagerman, R.J., Leehey, M., Heinrichs, W., Tassone, F., Wilson, R., Hills, J., Grigsby, J., Gage, B., and Hagerman, P.J. (2001). Intention tremor, parkinsonism, and generalized brain atrophy in male carriers of fragile X. Neurology 57, 127–130.

Hoffman, G.E., Le, W.W., Entezam, A., Otsuka, N., Tong, Z.B., Nelson, L., Flaws, J.A., McDonald, J.H., Jafar, S., and Usdin, K. (2012). Ovarian abnormalities in a mouse model of fragile X primary ovarian insufficiency. J Histochem Cytochem 60, 439–456.

Huang, J.Y., and Rosenwaks, Z. (2014). Assisted reproductive techniques. Methods Mol Biol 1154, 171–231.

Hunter, J., Rivero-Arias, O., Angelov, A., Kim, E., Fotheringham, I., and Leal, J. (2014). Epidemiology of fragile X syndrome: a systematic review and meta-analysis. Am J Med Genet A 164A, 1648–1658.

Hutchinson, A.P., Pereira, N., Lilienthal, D.P., Coveney, S., Lekovich, J.P., Elias, R.T., and Rosenwaks, Z. (2018). Impact of FMR1 Pre-Mutation Status on Blastocyst Development in Patients Undergoing Pre-Implantation Genetic Diagnosis. Gynecol Obstet Invest 83, 23–28.

Jo, Y.J., Jang, W.I., Namgoong, S., and Kim, N.H. (2015). Actin-capping proteins play essential roles in the asymmetric division of maturing mouse oocytes. J Cell Sci 128, 160–170.

Karen Nenonene, E., Trottier-Lavoie, M., Marchais, M., Bastien, A., Gilbert, I., Macaulay, A.D., Khandjian, E.W., Maria Luciano, A., Lodde, V., Viger, R.S., et al. (2023). Roles of the cumulus-oocyte transzonal network and the Fragile X protein family in oocyte competence. Reproduction 165, 209–219.

Kenneson, A., Zhang, F., Hagedorn, C.H., and Warren, S.T. (2001). Reduced FMRP and increased FMR1 transcription is proportionally associated with CGG repeat number in intermediate-length and premutation carriers. Hum Mol Genet 10, 1449–1454.

Kiehart, D.P., and Franke, J.D. (2002). Actin dynamics: the arp2/3 complex branches out. Curr Biol 12, R557–559.

Laggerbauer, B., Ostareck, D., Keidel, E.M., Ostareck-Lederer, A., and Fischer, U. (2001). Evidence that fragile X mental retardation protein is a negative regulator of translation. Hum Mol Genet 10, 329–338.

Lekovich, J., Man, L., Xu, K., Canon, C., Lilienthal, D., Stewart, J.D., Pereira, N., Rosenwaks, Z., and Gerhardt, J. (2018). CGG repeat length and AGG interruptions as indicators of fragile X-associated diminished ovarian reserve. Genet Med 20, 957–964.

Li, Z., Zhang, Y., Ku, L., Wilkinson, K.D., Warren, S.T., and Feng, Y. (2001). The fragile X mental retardation protein inhibits translation via interacting with mRNA. Nucleic Acids Res 29, 2276–2283.

Longo, F.J., and Chen, D.Y. (1985). Development of cortical polarity in mouse eggs: involvement of the meiotic apparatus. Dev Biol 107, 382–394.

Man, L., Lekovich, J., Rosenwaks, Z., and Gerhardt, J. (2017). Fragile X-Associated Diminished Ovarian Reserve and Primary Ovarian Insufficiency from Molecular Mechanisms to Clinical Manifestations. Front Mol Neurosci 10, 290.

Metchat, A., Eguren, M., Hossain, J.M., Politi, A.Z., Huet, S., and Ellenberg, J. (2015). An actin-dependent spindle position checkpoint ensures the asymmetric division in mouse oocytes. Nat Commun 6, 7784.

Michaelsen-Preusse, K., Zessin, S., Grigoryan, G., Scharkowski, F., Feuge, J., Remus, A., and Korte, M. (2016). Neuronal profilins in health and disease: Relevance for spine plasticity and Fragile X syndrome. Proc Natl Acad Sci U S A 113, 3365–3370.

Nikalayevich, E., Letort, G., de Labbey, G., Todisco, E., Shihabi, A., Turlier, H., Voituriez, R., Yahiatene, M., Pollet-Villard, X., Innocenti, M., et al. (2024). Aberrant cortex contractions impact mammalian oocyte quality. Dev Cell 59, 841–852 e847.

Pepling, M.E. (2006). From primordial germ cell to primordial follicle: mammalian female germ cell development. Genesis 44, 622–632.

Reeve, S.P., Bassetto, L., Genova, G.K., Kleyner, Y., Leyssen, M., Jackson, F.R., and Hassan, B.A. (2005). The Drosophila fragile X mental retardation protein controls actin dynamics by directly regulating profilin in the brain. Curr Biol 15, 1156–1163.

Rockwell, A.L., and Hongay, C.F. (2019). The m(6)A Dynamics of Profilin in Neurogenesis. Front Genet 10, 987.

Rosario, R., Stewart, H.L., Choudhury, N.R., Michlewski, G., Charlet-Berguerand, N., and Anderson, R.A. (2022). Evidence for a fragile X messenger ribonucleoprotein 1 (FMR1) mRNA gain-of-function toxicity mechanism contributing to the pathogenesis of fragile X-associated premature ovarian insufficiency. FASEB J 36, e22612.

Rotty, J.D., Wu, C., Haynes, E.M., Suarez, C., Winkelman, J.D., Johnson, H.E., Haugh, J.M., Kovar, D.R., and Bear, J.E. (2015). Profilin-1 serves as a gatekeeper for actin assembly by Arp2/3-dependent and -independent pathways. Dev Cell 32, 54–67.

Schuh, M., and Ellenberg, J. (2007). Self-organization of MTOCs replaces centrosome function during acentrosomal spindle assembly in live mouse oocytes. Cell 130, 484–498.

Seo, S.S., Louros, S.R., Anstey, N., Gonzalez-Lozano, M.A., Harper, C.B., Verity, N.C., Dando, O., Thomson, S.R., Darnell, J.C., Kind, P.C., et al. (2022). Excess ribosomal protein production unbalances translation in a model of Fragile X Syndrome. Nat Commun 13, 3236.

Sherman, S.L., Curnow, E.C., Easley, C.A., Jin, P., Hukema, R.K., Tejada, M.I., Willemsen, R., and Usdin, K. (2014). Use of model systems to understand the etiology of fragile X-associated primary ovarian insufficiency (FXPOI). J Neurodev Disord 6, 26.

Suarez, C., Carroll, R.T., Burke, T.A., Christensen, J.R., Bestul, A.J., Sees, J.A., James, M.L., Sirotkin, V., and Kovar, D.R. (2015). Profilin regulates F-actin network homeostasis by favoring formin over Arp2/3 complex. Dev Cell 32, 43–53.

Sun, S.C., Wang, Z.B., Xu, Y.N., Lee, S.E., Cui, X.S., and Kim, N.H. (2011). Arp2/3 complex regulates asymmetric division and cytokinesis in mouse oocytes. PLoS One 6, e18392.

Tassone, F., Beilina, A., Carosi, C., Albertosi, S., Bagni, C., Li, L., Glover, K., Bentley, D., and Hagerman, P.J. (2007). Elevated FMR1 mRNA in premutation carriers is due to increased transcription. RNA 13, 555–562.

Tassone, F., Hagerman, R.J., Taylor, A.K., Gane, L.W., Godfrey, T.E., and Hagerman, P.J. (2000). Elevated levels of FMR1 mRNA in carrier males: a new mechanism of involvement in the fragile-X syndrome. Am J Hum Genet 66, 6–15.

Telfer, E.E., Grosbois, J., Odey, Y.L., Rosario, R., and Anderson, R.A. (2023). Making a good egg: human oocyte health, aging, and in vitro development. Physiol Rev 103, 2623–2677.

Todd, P.K., Oh, S.Y., Krans, A., He, F., Sellier, C., Frazer, M., Renoux, A.J., Chen, K.C., Scaglione, K.M., Basrur, V., et al. (2013). CGG repeat-associated translation mediates neurodegeneration in fragile X tremor ataxia syndrome. Neuron 78, 440–455.

Verlhac, M.H., Lefebvre, C., Guillaud, P., Rassinier, P., and Maro, B. (2000). Asymmetric division in mouse oocytes: with or without Mos. Curr Biol 10, 1303–1306.

Wang, F., Zhang, L., Zhang, G.L., Wang, Z.B., Cui, X.S., Kim, N.H., and Sun, S.C. (2014). WASH complex regulates Arp2/3 complex for actin-based polar body extrusion in mouse oocytes. Sci Rep 4, 5596.

Wei, Z., Greaney, J., Zhou, C., and H, A.H. (2018). Cdk1 inactivation induces post-anaphase-onset spindle migration and membrane protrusion required for extreme asymmetry in mouse oocytes. Nat Commun 9, 4029.

Willemsen, R., Levenga, J., and Oostra, B.A. (2011). CGG repeat in the FMR1 gene: size matters. Clinical genetics 80, 214–225.

Yang, L., Duan, R., Chen, D., Wang, J., and Jin, P. (2007). Fragile X mental retardation protein modulates the fate of germline stem cells in Drosophila. Hum Mol Genet 16, 1814–1820.

